# The Pathogenic R5L Mutation Disrupts Formation of Tau Complexes on the Microtubule by Altering Local N-Terminal Structure

**DOI:** 10.1101/2021.10.15.464588

**Authors:** Alisa Cario, Adriana Savastano, Neil B. Wood, Zhu Liu, Michael J. Previs, Adam G. Hendricks, Markus Zweckstetter, Christopher L. Berger

## Abstract

The microtubule-associated protein (MAP) Tau is an intrinsically disordered protein (IDP) primarily expressed in axons, where it functions to regulate microtubule dynamics, modulate motor protein motility, and participate in signaling cascades. Tau misregulation and point mutations are linked to neurodegenerative diseases, including Progressive Supranuclear Palsy (PSP), Pick’s Disease and Alzheimer’s disease. Many disease-associated mutations in Tau occur in the C-terminal microtubule-binding domain of the protein. Effects of C-terminal mutations in Tau have led to the widely accepted disease-state theory that missense mutations in Tau reduce microtubule-binding affinity or increase Tau propensity to aggregate. Here, we investigate the effect of an N-terminal disease-associated mutation in Tau, R5L, on Tau-microtubule interactions using an *in vitro* reconstituted system. Contrary to the canonical disease-state theory, we determine the R5L mutation does not reduce Tau affinity for the microtubule using Total Internal Reflection Fluorescence (TIRF) Microscopy. Rather, the R5L mutation decreases the ability of Tau to form larger order complexes, or Tau patches, at high concentrations of Tau. Using Nuclear Magnetic Resonance (NMR), we show that the R5L mutation results in a local structural change that reduces interactions of the projection domain in the presence of microtubules. Altogether, these results challenge both the current paradigm of how mutations in Tau lead to disease and the role of the projection domain in modulating Tau behavior on the microtubule surface.

**Significance Statement:** The microtubule-associated protein Tau is strongly linked to a number of neurological diseases. Disease onset is typically associated with weakened interaction with the microtubule, but this widely accepted model is based on hyperphosphorylation or mutations within the C-terminal microtubule-binding domain of Tau. Here, we find an N-terminal disease-associated mutation in Tau, R5L, does not reduce Tau affinity for microtubules, but instead modifies the N-terminal structure, altering Tau’s behavior and ability to condense on the microtubule surface. Our findings challenge the current paradigms of both how mutations in Tau lead to disease and the functional role of the N-terminal region of Tau.

## Introduction

The microtubule-associated protein (MAP) Tau is an intrinsically disordered protein (IDP) highly expressed in the axon where it functions to regulate microtubule dynamics (1, 2), modulate motor protein motility (3–6), and participate in signaling cascades (7, 8). Misregulation and aggregation of Tau is linked to a number of neurodegenerative diseases known as Tauopathies. These include Alzheimer’s disease, Frontotemporal Dementia, and Progressive Supranuclear Palsy (PSP) (9–13). Disease-associated perturbations to Tau, including hyperphosphorylation, are generally thought to initiate the disease state by decreasing the affinity for microtubules (14–16). In support, there are over 50 disease-associated mutations in Tau and many have been shown to either reduce Tau affinity for the microtubule (17, 18) or promote Tau aggregation (18). However, many of the studied alterations are within the regions of Tau that directly bind to the microtubule, and less is understood about disease-associated mutations outside of the microtubule-binding region of Tau.

There are six isoforms of Tau expressed in the human brain (19), and although the exact composition can vary dependent on alternative splicing (20–22), all share the same functional domains. Three or four imperfect microtubule-binding repeats directly interface with the microtubule (23–28) in an extended conformation along a single protofilament (29), largely contributing to the high affinity of Tau for the microtubule. Adjacent to the microtubule binding repeats is the proline rich region which helps mediate microtubule binding (24, 27, 28, 30). The structurally flexible N-terminal region of Tau, known as the projection domain, also contains alternatively spliced acidic inserts (zero, one or two), and has the lowest contribution to the overall binding affinity of Tau for the microtubule (23, 31).

Additionally, Tau interaction with the microtubule is complicated by multiple binding modes. Tau binding is concentration dependent. At low concentrations, Tau binds to the microtubule in a dynamic equilibrium between static and diffusive binding states (32, 33). However, as the concentration of Tau increases, Tau binds nonuniformly and forms larger order complexes, referred to as Tau patches (6), Tau cohesive islands (34), or Tau condensates (35). Tau complex formation on the microtubule is not restricted to *in vitro* studies as microtubule-bound complexes have been seen in neurons and other cell types (35–37). Interestingly, in addition to the clear role for the C-terminal microtubule-binding region of Tau in complex formation, the N-terminal projection domain has been shown to be necessary in Tau cohesive island formation (34).

Although many of the mutations in Tau are found within the C-terminal half the protein, disease-associated mutations also occur in N-terminal projection domain, specifically the R5 mutations (R5L, R5H), associated with PSP (38). The R5L mutation has been shown to reduce microtubule assembly, although the mechanism behind this change is unclear (39, 40). Deletion of the projection domain does not reduce Tau affinity for the microtubule (26, 41, 42), but is necessary for Tau complex formation (34). Thus, we hypothesize that the N-terminal disease-associated mutation R5L alters Tau binding behavior on the microtubule, disrupting patch formation without altering Tau affinity. Our results suggest the R5L mutation is unlike other disease-associated mutations and alters Tau-microtubule interactions in a previously unreported manner, leading insight into a new potential disease mechanism.

## Results

### R5L-Tau reduces occupancy but does not alter affinity on Taxol-microtubules

To study the effect of R5L-Tau on Tau:microtubule interactions, the R5L mutation was cloned into the 3RS isoform of Tau and the affinity of R5L-Tau was assessed relative to WT-Tau using a TIRF (Total Internal Reflection Fluorescence) Microscopy binding assay as previously reported (43). Fluorescently labeled Tau (10-300 nM WT-Tau or R5L-Tau) was incubated with microtubules stabilized with paclitaxel (Taxol-microtubules) (Fig. 1A). The average Tau intensity on the microtubule was plotted against the Tau concentration and fit to a cooperative binding model (Fig. 1B). It was determined the R5L mutation did not significantly alter the affinity of Tau for Taxol-microtubules (WT-Tau K_D_ = 140 ± 19 nM, R5L-Tau K_D_ = 98 ± 10 nM) (Fig. 1B).

**Fig. 1.**
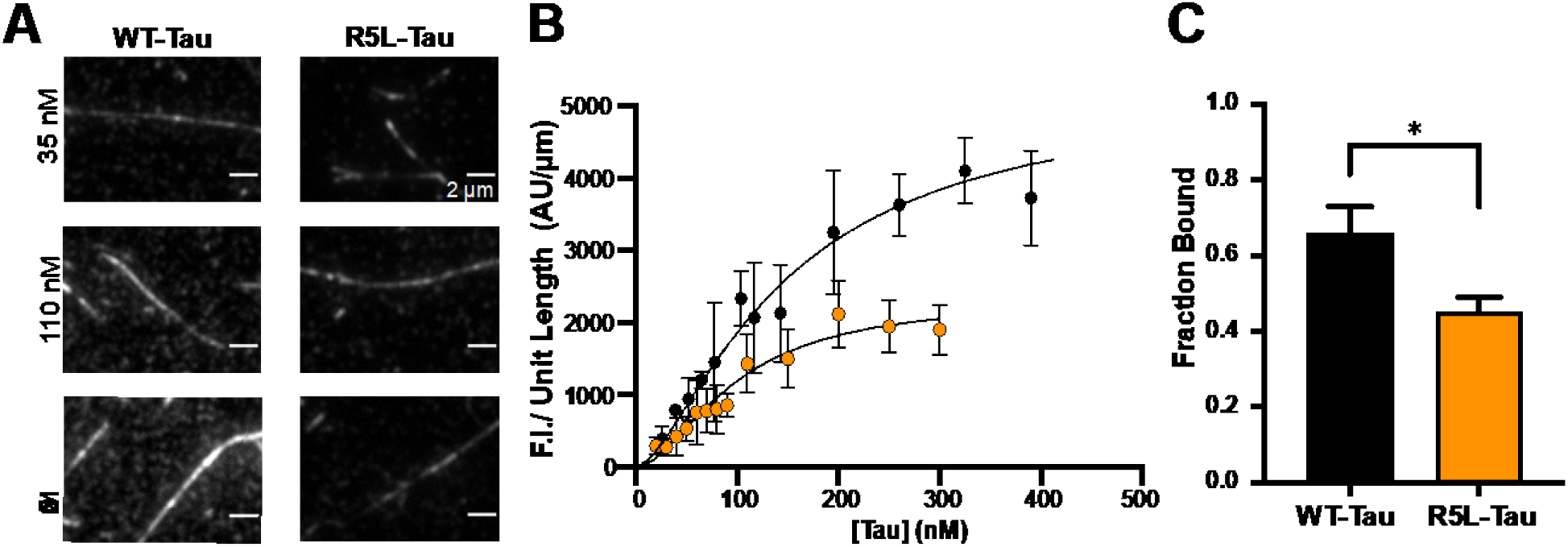
The R5L mutation reduces Tau occupancy on Taxol-microtubules. **A.** Representative images of WT-Tau (left) and R5L-Tau (right) decoration of Taxol-microtubules at varying Tau concentrations. **B.** TIRF based binding assays fit to a cooperative binding model comparing WT-Tau (black, K_D_ = 140 ± 19 nM with Hill Coefficient = 1.4 ± 0.1) and R5L-Tau (orange, K_D_ = 98 ± 10 nM with Hill Coefficient = 1.8 ± 0.2). Data are mean ± 95% CI. (N = 4) **C.** Microtubule pelleting assay comparing relative fluorescence bound Tau of WT-Tau (black, 0.66 ± 0.07) or R5L-Tau (orange, 0.45 ± 0.04) at 100 nM Tau and 1 μM Taxol-microtubules. Data are mean ± SD (N = 3). Statistical analysis was performed using students t-test (* p < 0.05).

Interestingly, the TIRF binding assay revealed a two-fold difference in fluorescence intensity between WT-Tau and R5L-Tau at binding saturation, despite similar affinities (Fig. 1B). It was determined this was not due to a difference in labeling efficiency or quenching of the fluorescent probe (Fig. S9). Labeling efficiency was determined by either a Modified Lowry assay in conjunction with absorbance to measure dye concentration or through Liquid Chromatography-mass spectrometry (LCMS) to determine the percent of modified cysteine residues (Supplemental methods). Therefore, we hypothesized the R5L mutation reduces the occupancy of Tau on microtubules without altering affinity, leading to less Tau bound to microtubules at binding saturation. To test this hypothesis, a microtubule pelleting assay was performed by incubating 100 nM labeled WT-Tau or R5L-Tau with varying concentrations of Taxol-microtubules (50 nM – 1 μM). The relative amount of Tau bound to Taxol-microtubules was assessed by measuring the fluorescence in the supernatant relative to the pellet. At saturating conditions, less R5L-Tau bound to microtubules compared to WT-Tau (Fig. 1C).

### The R5L mutation disrupts Tau patch formation

The TIRF binding assays also showed non-uniform binding of Tau along the microtubule as the concentration increased (Fig. 1A, Movie S1-S2). This is consistent with other reports of Tau forming larger order complexes on the microtubule surface, known as Tau patches, cohesive islands or condensates (6, 32, 33, 35). Therefore, we studied the effect of the R5L mutation on Tau patches by incubating 250 nM Tau (WT-Tau or R5L-Tau) with Taxol-microtubules using TIRF microscopy (Fig. 2A). Patch frequency was quantified as the number of patches per unit length along the microtubule. WT-Tau formed twice as many patches per micron compared to R5L-Tau (WT-Tau = 0.36 ± 0.21, R5L-Tau = 0.18 ± 0.21) (Fig. 2B). WT-Tau patches also contained approximately 50% more molecules than R5L-Tau patches (WT-Tau = 3.3 ± 1.1 molecules/patch, R5L-Tau = 2.1 ± 1.8 molecules/patch) (Fig. 2C). Furthermore, we examined the fluorescence intensity distribution along the microtubule (Fig. 2D). There was a higher mean fluorescence intensity and a broader distribution for WT-Tau compared to R5L-Tau. Thus, WT-Tau forms more numerous and larger patches compared to R5L-Tau.

**Fig. 2.**
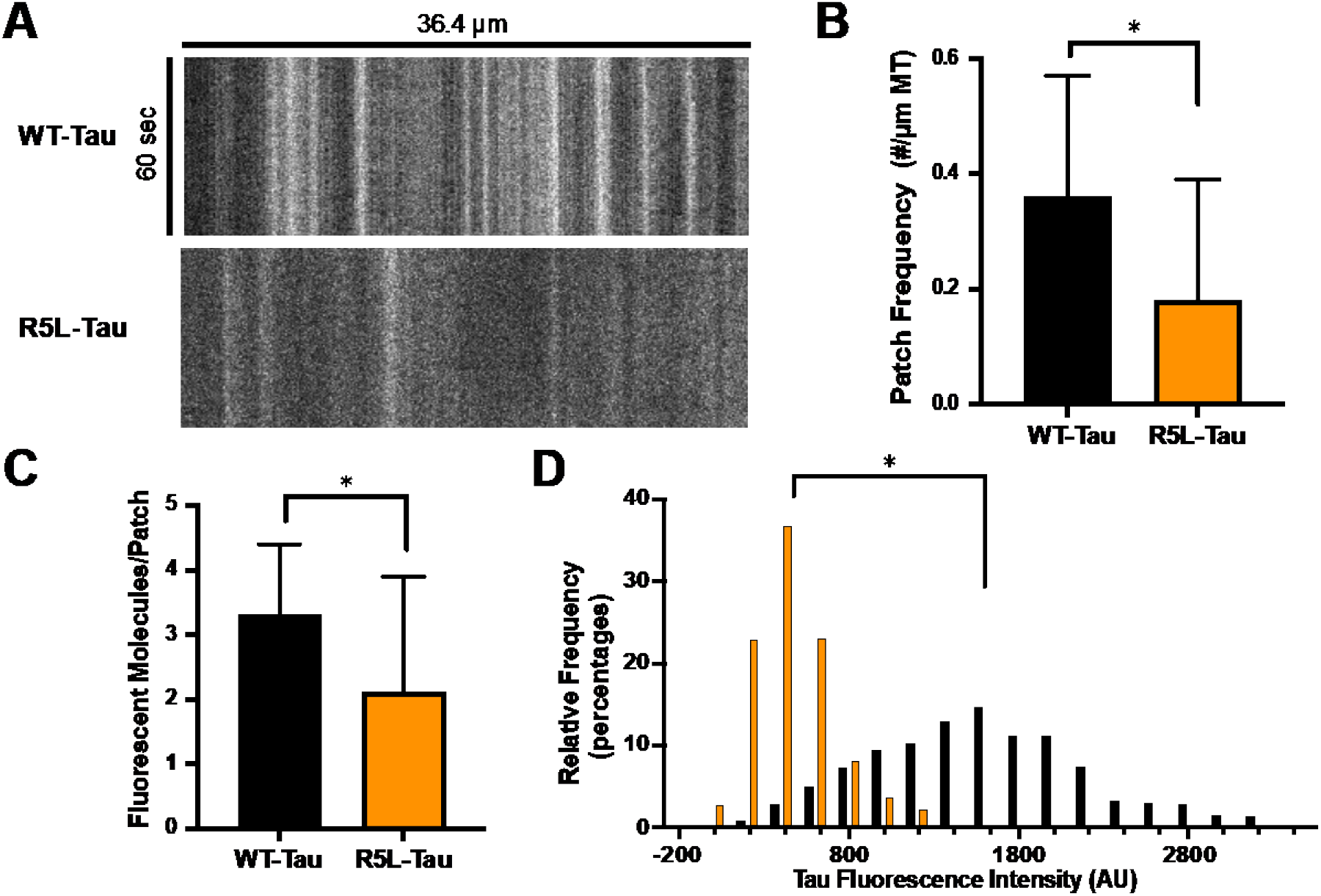
The R5L mutation alters Tau patches on Taxol-microtubules. Tau patches were studied using 250 nM Alexa 488 Tau on Taxol-microtubules. **A.** Representative kymographs of WT-Tau (top) and R5L-Tau (bottom) at 250 nM Tau concentration. **B.** Patch frequency of WT-Tau (black, 0.36 ± 0.21 patches/μm) and R5L-Tau (orange, 0.18 ± 0.21 patches/μm). Data are mean ± SD (N = 80 kymographs). Statistical analysis was performed using students t-test (*p < 0.05). **C**. Number of fluorescent molecules per patch of WT-Tau (black, 3.3 ± 1.1 fluorescent molecules/patch) and R5L-Tau (orange, 2.1 ± 1.8 fluorescent molecules/patch). Data are mean ± SD (N = 80 kymographs). Statistical analysis was performed using students t-test (*p < 0.05). **D.** Histograms of normalized WT-Tau (black) and R5L-Tau (orange) Tau fluorescence intensity on microtubules. Values above 1% are shown. (N = 80 kymographs). Statistical analysis was performed using Mann-Whitney test (*p < 0.01).

### R5L-Tau shifts binding behavior at high concentrations on Taxol-microtubules

We investigated the difference in the binding behavior of individual R5L-Tau and WT-Tau molecules on Taxol-microtubules using TIRF microscopy. Previous work has shown that at low concentrations of Tau, Tau binds to the microtubule in a dynamic equilibrium between static and diffusive binding states (32, 33). (33). Therefore, we studied the binding behavior of 500 pM WT-Tau or R5L-Tau bound to Taxol-microtubules and measured the binding state equilibrium, diffusion coefficient and dwell times of both static and diffusive events (representative kymographs are shown in Fig. S1). There was no significant difference in the binding state equilibrium at low concentrations of both WT-Tau and R5L-Tau, with approximately half of the binding events observed in the static state. (Table 1, Fig. S2). Additionally, the R5L mutation had little effect on the diffusion coefficient or dwell time of individual molecules in either the static or diffusive states (Table 1, Fig. S3-S4). Overall, this data showed little difference in the binding behavior between WT-Tau and R5L-Tau at low concentrations.

**Table 1.** Summary of WT-Tau and R5L-Tau behavior on microtubules. Low concentration studies were performed with 500 pM Alexa 647 labeled Tau on microtubules. For Taxol-microtubules, high concentration studies were performed through a spiking experiment, incubating 250 nM Alexa 488 labeled Tau with 300 pM Alexa 647 labeled Tau. For GMPCPP microtubules, high concentration studies were performed through a spiking experiment, incubating 400 nM Alexa 488 Tau with 300 pM Alexa 647 Tau. Dwell times and diffusion coefficients are represented as median ± 95%CI. (*p < 0.01) relative to WT-Tau under same conditions.

|  |  |  | Percent<br>static | Diffusion<br>Coefficient | Dwell time<br>static | Dwell time<br>diffusive |
| --- | --- | --- | --- | --- | --- | --- |
| | | | % | $\mu m^2/sec$ | sec | sec |
| **Taxol** | Low Conc. | WT-Tau | 45 | $0.29 \pm 0.04$ | $1.5 \pm 0.3$ | $1.4 \pm 0.1$ |
| | | R5L-Tau | 57 | $0.33 \pm 0.04$ | $1.6 \pm 0.2$ | $1.2 \pm 0.2$ |
| | High Conc. | WT-Tau | 48 | $0.14 \pm 0.02$ | $1.8 \pm 0.4$ | $2.4 \pm 0.3$ |
| | | R5L-Tau | 17* | $0.21 \pm 0.02^*$ | $2.1 \pm 0.3$ | $2.3 \pm 0.2$ |
| **GMPCPP** | Low Conc. | WT-Tau | 26 | $0.75 \pm 0.10$ | $2.2 \pm 0.5$ | $1.1 \pm 0.2$ |
| | | R5L-Tau | 23 | $0.75 \pm 0.10$ | $1.0 \pm 0.2^*$ | $0.9 \pm 0.1$ |
| | High Conc. | WT-Tau | 15 | $0.50 \pm 0.08$ | $2.0 \pm 0.6$ | $1.5 \pm 0.4$ |
| | | R5L-Tau | 14 | $0.44 \pm 0.09$ | $1.1 \pm 0.1$ | $1.2 \pm 0.1$ |

However, at higher Tau concentrations, complexes comprised of statically bound Tau molecules form on the microtubule (35). Due to the differences in R5L-Tau patch formation, we examined the binding behavior of individual Tau molecules at high concentration by performing a spiking experiment where 250 nM Alexa 488 WT- or R5L-Tau was spiked with 300 pM Alexa 647 Tau (Movie S3-S4). We examined the static-diffusive equilibrium, diffusion coefficient and the dwell times. At high concentrations of Tau, 48% of WT-Tau bound statically compared to 17% for R5L-Tau (Table 1, Fig. S2). When bound diffusively, WT-Tau had a lower diffusion coefficient compared to R5L-Tau (WT-Tau = 0.14 ± 0.02 μm^2^/sec, R5L-Tau =0.21 ± 0.02 μm^2^/sec) (Table 1, Fig. S3). There was little difference in the dwell times of either static or diffusive molecules (Table 1, Fig. S4). Thus, at high concentration R5L-Tau shifts to a predominantly diffusive state (Table 1).

### R5L-Tau reduces occupancy but does not alter affinity on GMPCPP-microtubules

On Taxol-microtubules, the R5L mutation had two effects: less Tau bound to the microtubule and fewer Tau patches. However, it was unclear if these two effects of the R5L mutation were linked. Previous work from our lab has shown that in comparison to Taxol-microtubules, Tau binds as single molecules and does not form complexes on microtubules stabilized with guanosine 5’-[(α,β)-methyleno] triphosphate sodium salt (GMPCPP-microtubules) (33). Consistent with this data, recent work shows Tau condensates do not form on GMPCPP-microtubules (35). Therefore, we studied R5L-Tau binding to GMPCPP microtubules to examine the effect of the R5L mutation independent of patch formation. The TIRF binding assay indicated that, as expected, both WT-Tau and R5L-Tau had a lower affinity for GMPCPP microtubules compared to Taxol-microtubules (44, 45). However, there was little difference in affinity between WT-Tau and R5L-Tau (WT-Tau K_D_ = 300 ± 25 nM, R5L-Tau K_D_ = 209 ± 20 nM) (Fig. 3B, Movies S5-S6). Interestingly, there was a two-fold change in fluorescence intensity between WT-Tau and R5L-Tau at saturating concentrations (Fig. 3B). Microtubule pelleting assays confirmed that less R5L-Tau bound to GMPCPP-microtubules compared to WT-Tau (Fig. 3C).

**Fig. 3.**
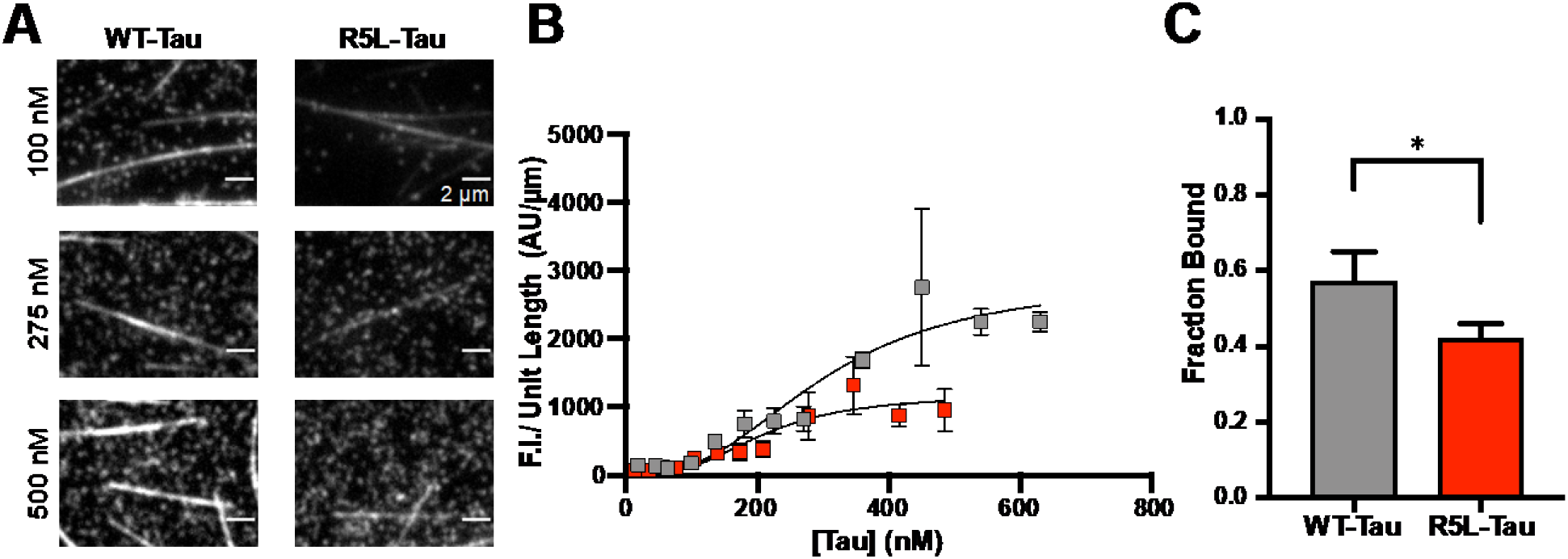
The R5L mutation reduces occupancy on GMPCPP-microtubules. **A.** Representative images of WT-Tau (left) and R5L-Tau (right) decoration of GMPCPP-microtubules at varying Tau concentrations. **B.** TIRF based binding assays comparing WT-Tau (grey, K_D_ = 300 ± 25 nM with Hill Coefficient = 2.6 ± 0.3) and R5L-Tau (red, 209 ± 20 nM with a Hill Coefficient = 2.9 ± 0.4). Data are mean ± 95% CI (N = 4) **C.** Microtubule pelleting assay comparing relative fluorescence bound Tau of 300 nM WT-Tau (grey, 0.57 ± 0.08) or R5L-Tau (red, 0.42 ± 0.04) bound to 1 μM GMPCPP-microtubules. Data are mean ± SD (N = 3) Statistical analysis was performed using students t-test (*p < 0.05).

### The R5L mutation does not alter binding behavior on GMPCPP-microtubules

To further understand the effect of R5L-Tau on GMPCPP-microtubules, we studied the binding behavior at low concentrations of Tau (500 pM WT-Tau or R5L-Tau) (Table 1). At low concentrations, both WT-Tau and R5L-Tau shift towards diffusive binding on GMPCPP-microtubules, consistent with previous work from our lab (33), but there was no significant difference in the overall binding behavior of WT-Tau vs. R5L-Tau. 26%of WT-Tau bound in the static state compared to 23% R5L-Tau (Table 1, Fig. S6), and there was no difference in diffusion coefficient or diffusive dwell time. However, when bound statically, WT-Tau had a higher dwell time compared to R5L-Tau (WT-Tau = 2.7 ± 0.4 seconds, R5L-Tau = 1.9 ± 0.3 seconds) (Table 1, Figs. S7-S8).

A shift in binding behavior for R5L-Tau occurred at high concentrations on Taxol-microtubules (Table 1, Fig. S2). Therefore, we studied the binding behavior at high concentrations of Tau on GMPCPP-microtubules by performing a spiking experiment where we incubated 400 nM Alexa 488 WT- or R5L-Tau spiked with 300 pM Alexa 647 Tau to visualize individual molecules on GMPCPP-microtubules (Table 1; representative kymographs are shown in Fig. S5). At high concentrations, 15% of WT-Tau bound statically compared to 14% R5L-Tau (Table 1, Fig. S6). Similar to low concentrations of Tau, there was little difference in the diffusion coefficient or diffusive dwell time (Table 1, Figs. S7-S8). However, WT-Tau had a higher dwell time compared to R5L-Tau (WT-Tau = 2.8 ±1.5 seconds, R5L-Tau = 1.9 ± 0.9 seconds) in the static state (Table 1, Fig. S8), albeit this a small percentage of the total binding events. Contrary to Taxol-microtubules, we did not see any significant differences in binding behavior on GMPCPP-microtubules at high concentrations of Tau.

### The R5L mutation alters the local structure of Tau

We then examined the impact of the R5L mutation on the secondary structure and global conformation of Tau using circular dichroism (CD) and dynamic light scattering (DLS). Both WT-Tau and R5L-Tau had similar CD spectra, typical for random coil conformation with minima at ~200 nm (Fig. 4A) (46), and hydrodynamic radii (R_H_) as determined by DLS (Fig. 4A). Two dimensional ^1^H-^15^N heteronuclear single quantum coherence (HSQC) experiments Nuclear Magnetic Resonance (NMR) spectroscopy of 10 μM ^15^N-labelled Tau (WT-Tau or R5L-Tau) was then used to obtain residue-specific structural information (47, 48) on the effect of the R5L mutation. Alterations in chemical shift were observed within 5-10 residues of the site of mutation, suggesting a local structural change restricted to the first 20 amino acids within the N-terminal projection domain (Fig. 4B-D).

**Fig. 4.**
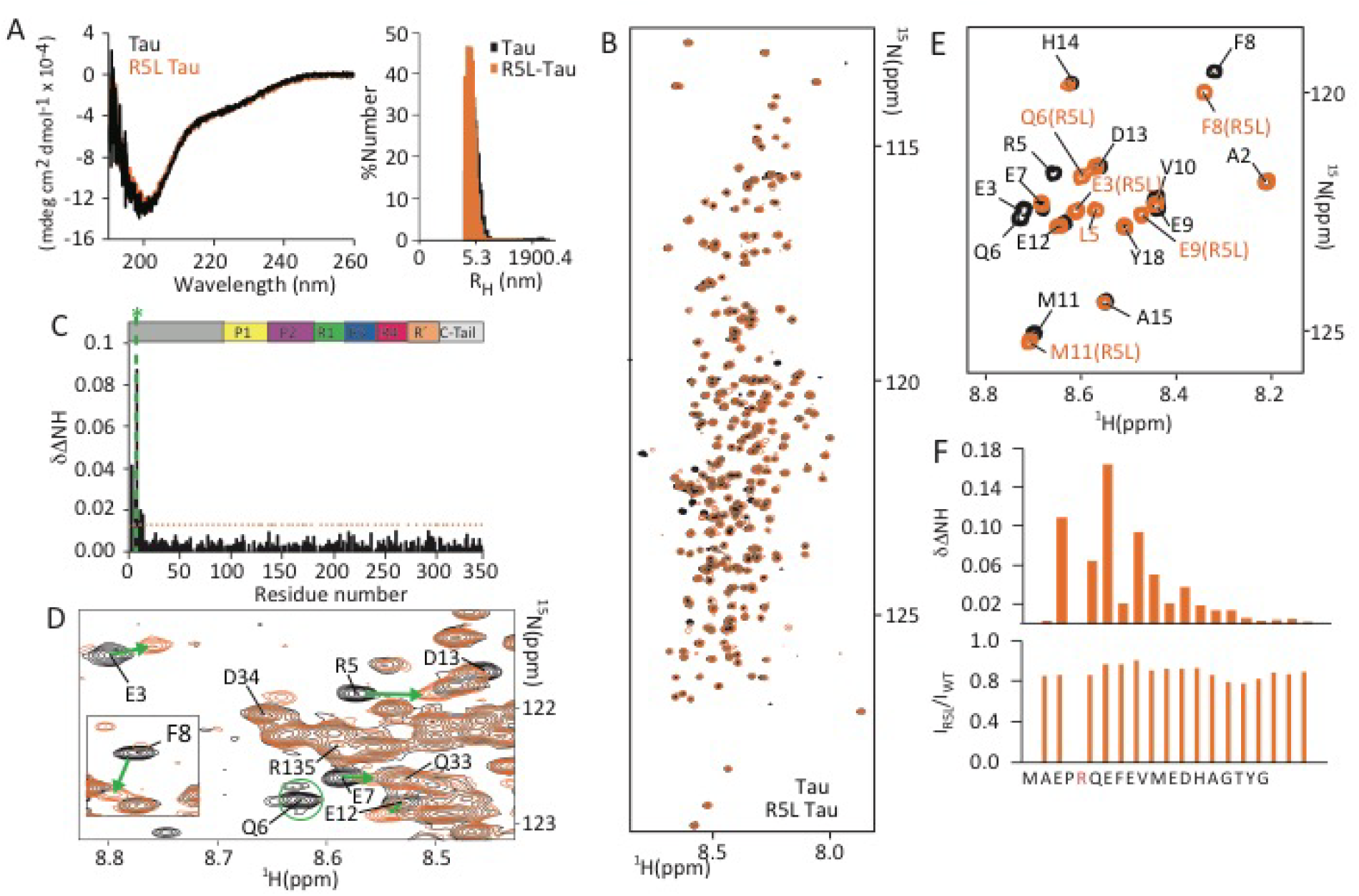
Changes in the structural dynamics near the R5L mutation of Tau. **A.** Structural impact of the R5L mutation probed by circular dichroism (CD; left) and dynamic light scattering (DLS; right). CD data are reported as Mean Residue Ellipticity (MRE) for 10 μM samples of WT-Tau and R5L-Tau. **B.** Superposition of 2D ^1^H^15^N-HSQC spectra of WT-Tau (black) and R5L-Tau (orange). **C.** Residue-specific ^1^H^15^N chemical shift perturbation of Tau upon mutation. The N-terminal region close to the mutated site (indicated by a dashed, green line and the asterisk) shows chemical shift perturbations above the threshold (dashed orange line). The threshold was calculated as the standard deviation between all the dDNH values and multiplied by a factor of 2. The domain organization of Tau is displayed on top of the plot. **D.** Zoom of selected regions of superimposed ^1^H-^15^N HSQC spectra of WT-Tau (black) and R5L-Tau (orange); shifts are indicated by green arrows. **E-F.** NMR characterization of WT-Tau and R5L-Tau peptides (residues 1-20 of Tau and R5L-Tau, respectively). Superposition of 2D ^1^H^15^N-HSQC spectra of Tau (black) and R5L-Tau (orange) peptides (E). Sequence-specific resonance assignments are indicated. Chemical shift perturbations (top panel) and intensity ratios (bottom panel) in the R5L-Tau peptide (F).

NMR spectra of IDPs like Tau are prone to signal overlap (48). We therefore further investigated the consequences of the R5L mutation using short peptides comprising the first 20 residues (WT-Tau or R5L-Tau (Fig. 4E-F). Similar to the full-length protein, chemical shift perturbations from natural abundance 2D ^1^H-^15^N HSQC experiments on 1 mM peptides were observed in direct proximity to the site of mutation (Fig. 4E). The two glutamate residues E3 and E7, which are one residue away from the R5L mutation, showed the largest perturbations (Fig. 4F). The combined analysis demonstrates that the structural changes caused by the R5L mutation are highly local and do not affect the microtubule-binding domain.

### The R5L mutation causes a reduction in projection domain interactions with the microtubule

To investigate the effect of the R5L mutation on the interaction of Tau with microtubules, we recorded 2D ^1^H-^15^N HSQC experiments of 10 μM ^15^N-labelled WT-Tau or R5L-Tau in the absence or presence of 20 μM Taxol-microtubules (Fig. 5A). Upon binding to microtubules the signal intensity of Tau residues interacting with the microtubule surface decreases (25), which can be visualized in an intensity ratio plot (Fig. 5A). The strongest drop in signal intensity is seen for both proteins in the second proline-rich region (P2) through the microtubule-binding repeats, which together form the microtubule-binding domain of Tau. In agreement with the highly local nature of the structural perturbations of the R5L mutation (Fig. 4C-F), the R5L mutation does not influence the high-affinity C-terminal microtubule-binding domain of Tau. In contrast, the R5L-mutation affects the N-terminal projection domain of Tau. The signal attenuation at the N-terminus of R5L-Tau is weaker when compared to WT-Tau (Fig. 5A), which becomes particularly evident upon inspection of the intensity difference plot (Fig. 5A). Notably, while the R5L-induced chemical shift perturbation was restricted to the N-terminal 15 residues in solution (Fig. 4C), the mutation affects up to 50 residues of the projection domain of Tau in the presence of microtubules.

**Fig. 5.**
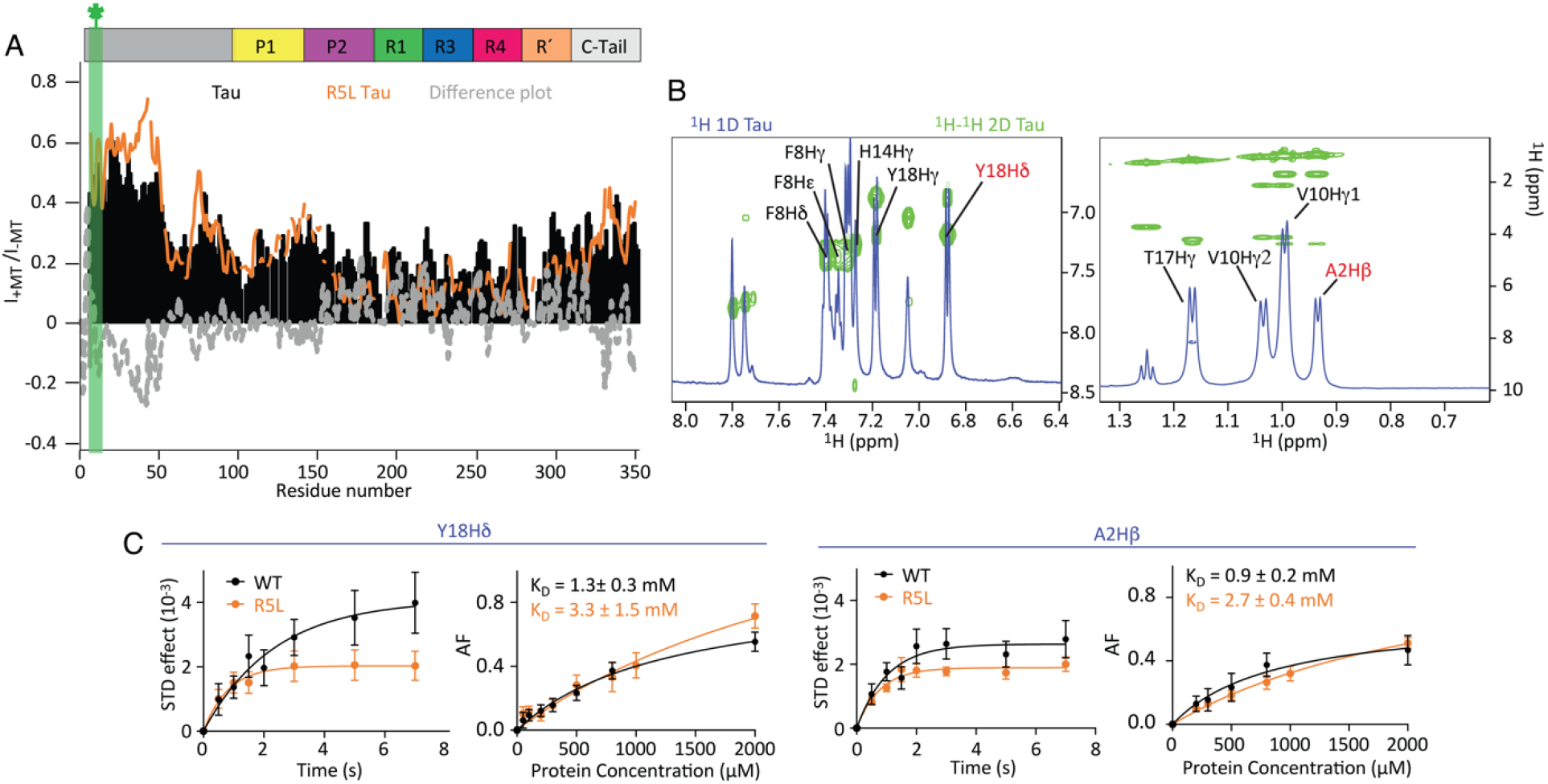
The R5L mutation attenuates the interaction of the projection domain of Tau in the presence of microtubule. **A.** Site-specific changes in the NMR signal intensities of WT-Tau (10 μM; black bars) or R5L-Tau (10 μM; orange line) upon addition of 20 μM Taxol-microtubules (1:2 molar ratio of Tau:tubulin heterodimer). I-MT and I+MT are the intensities of ^1^H-^15^N HSQC cross peaks in the absence and presence of microtubules, respectively. The dashed grey line displays the residue-specific differences between WT-Tau and R5L-Tau. The site of mutation is indicated by the green bar and asterisk. **B.** Superposition of the aromatic (left panel) and methyl (right panel) regions of ^1^H 1D and ^1^H -^1^H 2D TOCSY spectra of the Tau peptide (residues 1-20 of Tau) showing its assignment. The signals of Y18Hd and A2Hb (shown in red) were used for further STD NMR analysis. **C.** STD effect and K_D_ values calculated for Y18Hd and A2Hb of the N-terminal peptides of WT-Tau (black) and R5L-Tau (orange) for binding to 5 μM Taxol-microtubules.

To quantify the effect of the mutation on Tau binding to microtubules, we used Saturation Transfer Difference (STD)-NMR spectroscopy, which provides information on the residues involved in the binding with a larger protein (i.e. microtubules) (49–51). The efficiency of saturation transfer was investigated for the WT-Tau and R5L-Tau peptides in the presence of 5 μM Taxol-microtubules through the measurement of the STD-NMR signals at increasing saturation time. For selected protons, i.e. Y18Hδ and A2Hβ the STD build up curve is displayed (Fig. 5B). The analysis demonstrates that both peptides receive magnetization from the microtubules, but the saturation transfer is more efficient for the WT-Tau peptide. Subsequently, we determined the microtubule-binding affinity of the two peptides by measuring STD NMR experiments with fixed concentration of Taxol-microtubules and gradually increasing peptide concentrations (Fig. 5C). The K_D_ values obtained for Y18Hδ and A2Hβ of the WT-Tau peptide were 1.3 ± 0.3 mM and 0.9 ± 0.2 mM respectively. In case of the R5L-Tau peptide, we derived K_D_ values of 3.3 ± 1.5 mM and 2.7 ± 0.4 mM, respectively.

## Discussion

Missense mutations or hyperphosphorylation of Tau that lead to neurodegeneration, such as PSP, are generally thought to be initiated by a decrease in Tau’s interaction with the microtubule. A reduction in binding affinity has been demonstrated in previously studied disease-associated mutations in the microtubule binding region (e.g., Δ280K, P301L and V337M) (17, 52–56). However, the present work challenges the generally accepted model of Tauopathies as the R5L mutation, located in the N-terminal projection domain, does not alter Tau affinity for the microtubule. This is consistent with past studies where deletion of the entire projection domain does not reduce Tau affinity (26, 41, 42). In support of this finding, microtubule induced NMR signal broadening measurements show the largest attenuation of signal occurs within the microtubule binding region, with no differences observed between WT-Tau and R5L-Tau. Rather, R5L-Tau has a loss of microtubule-induced attenuation in the projection domain relative to WT-Tau, suggesting a decreased engagement of the projection domain of R5L-Tau in intermolecular or intramolecular interactions.

Although Tau is an intrinsically disordered protein, it populates an ensemble of conformations in solution where the N-terminal domain interacts with the C-terminal domain (31, 48, 57, 58). Measurements of the conformational ensemble of R5L-Tau through CD, DLS and NMR indicate no global conformational changes, consistent with no change in affinity, relative to WT-Tau. Alterations in the N-terminus that reduce Tau affinity such as phosphorylation of Y18 (43) cause global conformational changes to Tau (7), likely through disruption of these transient long range interactions. However, this is not seen with R5L-Tau as the R5L mutation only perturbs the local structure of Tau proximal to the mutation (up to ~10 AA from R5). Furthermore, no long-range effects are measured in the presence of microtubules, where changes to the microtubule-induced signal broadening remain local but are propagated approximately 50 residues from the site of the mutation.

However, the local changes in R5L-Tau alter Tau behavior and disrupts Tau patch formation on Taxol-microtubules. Tau is known form complexes on Taxol-microtubules which have been referred to as Tau patches (6), condensates (35), or cohesive islands (34). Similarly, microtubule bound Tau complexes have been shown to occur in neurons and other cell types (35–37). Although it is not clear whether these complexes are identical as experimental conditions differ, all Tau complexes share similar properties. Our previous work has shown that static Tau forms small complexes on Taxol-microtubules (33). Moreover, recent work has shown Tau condensates/cohesive islands are comprised of statically bound molecules (34, 35). Here, TIRF mobility assays show a concentration dependent shift in the binding behavior of R5L-Tau on Taxol-microtubules, indicating the R5L mutation disrupts the ability of Tau to form patches. At low concentrations, Tau patches do not form and there is no difference in the binding behavior between WT-Tau and R5L-Tau. The difference arises at higher Tau concentrations, where R5L-Tau shifts towards a diffusive state, in agreement with less Tau patch formation (Table 1). Further, R5L-Tau has a lower patch frequency and fewer molecules per patch compared to WT-Tau at high concentrations (Fig. 2B-D). Our NMR data reveals the R5L mutation causes changes in the local structure of the projection domain, consistent with previous work that showed the projection domain is necessary for formation of Tau cohesive islands (34), and phase separation properties of Tau condensates (35).

The ability of the R5L mutation to disrupt patch formation is further supported by our work on GMPCPP-microtubules, which do not form Tau complexes (33, 35). There is no change in binding behavior between WT-Tau and R5L-Tau on GMPCPP-microtubules, supporting the shift in binding behavior with the R5L mutation on Taxol-microtubules is due to disruption of patch formation (Table 1). Interestingly, there is an overall reduction in the occupancy of Tau on GMPCPP-microtubules compared to Taxol-microtubules, suggesting a shift towards diffusive binding contributes to the reduced occupancy of R5L-Tau on Taxol-microtubules. However, the R5L mutation does not alter the binding state equilibrium and yet reduces the occupancy on GMPCPP-microtubules, indicating the reduction in occupancy with the R5L mutation is more complicated than a shift in binding behavior, and likely due to the local structural changes within the projection domain.

In our experiments, the percent of static molecules for WT-Tau is lower than previously reported, specifically at the higher Tau concentrations (33, 43). However, binding state classification is complicated by the ability of a single event to contain both static and diffusive binding states (33, 43). Additionally, the binding state analysis used in the current study differs from our previous work and is more sensitive to diffusive binding. Further analysis of the classification of binding state indicates WT-Tau has a stronger dependence on the classification parameter at high Tau concentrations compared to low Tau concentrations, suggesting there are more binding events that contain both static and diffusive behavior at high Tau concentration (Fig. S10). Furthermore, the reduction in diffusion coefficient at high Tau concentrations is also consistent with events that exhibit a mix of static and diffusive binding (Table 1). Thus, events that contain both binding states are more likely to be classified as diffusive, decreasing the percent static molecules at higher Tau concentrations. We find that compared to R5L-Tau, WT-Tau has more static molecules at any classification parameter value (Fig. S10), further supporting the shift towards diffusive binding for R5L-Tau and disruption of patch formation.

Previously, we showed that arginine residues are important in the formation of salt bridges between Tau and tubulin (59). We hypothesize the mechanism by which the R5L mutation alters the behavior of microtubule-bound Tau is through loss of a salt bridge involving R5, thus altering the ensemble structural conformations occupied by the projection domain. However, due to the high flexibility of the projection domain and corresponding lack of stable structure, the specific salt bridge the R5L mutation disrupts is uncertain. The R5 residue could be engaged in a salt bridge within Tau, with other Tau molecules, or with the microtubule. These three models are not mutually exclusive and based on the dynamic nature of Tau, a loss of multiple transient interactions could contribute to the overall effect of the R5L mutation, which is to disrupt microtubule bound Tau patches (Fig. 6).

**Fig. 6.**
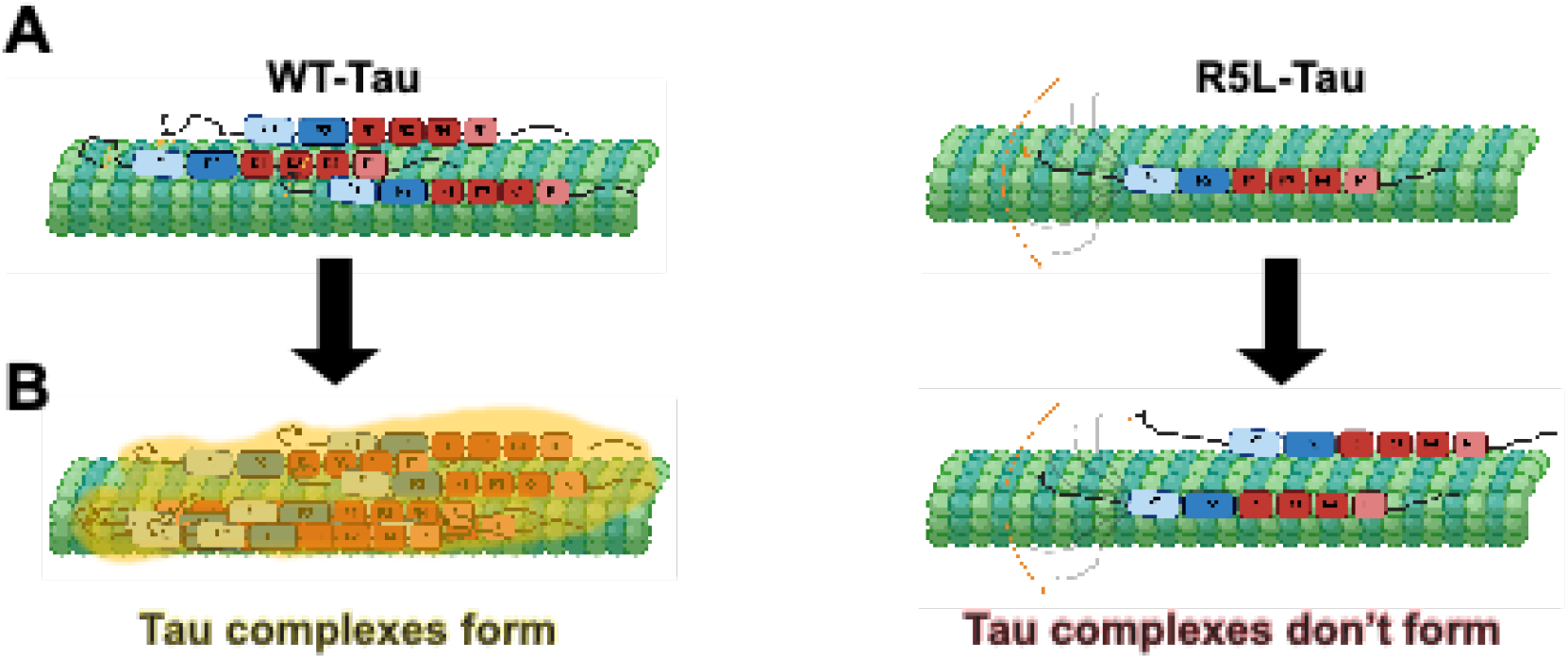
Mechanistic model for the impact of the R5L mutation on Tau binding to microtubules. The R5L mutation disrupts Tau condensation on the microtubule. **A.** The R5 residue is important for salt bridge formation in WT-Tau (left) that does not occur in R5L-Tau (right). **B.** The salt bridge contacts in WT-Tau (left) allow Tau condensation that does not occur in R5L-Tau (right).

The N-terminal region of Tau is largely acidic and the R5 residue could engage in an intramolecular salt bridge with a number of amino acids found within the projection domain. NMR data reveal the R5L mutation disrupts the local structure of Tau in solution, which could be attributed to the loss of an intramolecular salt bridge. However, the structural change is enhanced in the presence of microtubules and therefore, a loss of an intramolecular salt bridge alone does not fully explain the effect of the R5L mutation.

An additional possibility is the R5 residue is engaged in an intermolecular Tau salt bridge that is disrupted by the R5L mutation. The N-terminal domain of Tau has been shown to be important for intermolecular Tau interactions (60) and the possible formation of an “electrostatic zipper” between Tau molecules (61). An intermolecular Tau salt bridge could occur between highly acidic projection domains or with the C-terminal binding region of another Tau molecule, although the overall structure of Tau within a microtubule-induced complex remains unclear. Furthermore, the R5L mutation disrupts microtubule bound Tau complexes, which are suggested to be mediated by intermolecular Tau interactions involving the projection domain (34).

Finally, due to the largely acidic surface of the microtubule, it is feasible R5 is engaged in a salt bridge with tubulin. Microtubule induced NMR signal broadening show a decrease in R5L-Tau N-terminal interactions in the presence of microtubules, suggesting R5 transiently associates with the microtubule surface. Furthermore, STD measurements of peptides show a reduced affinity with the R5L mutation. Taken together, these experiments show the R5L mutation weakens the interaction of the projection domain with the microtubule. Our data is consistent with possible interactions of the projection and microtubule-binding domains on the same microtubule or an adjacent microtubule (62), depending on microtubule organization. For example, the R5 residue could interact with adjacent microtubules in densely packed bundles in the axon.

Here we find the main effect of the R5L mutation is disruption of Tau patches, suggesting an important physiological role for microtubule bound complex formation. Tau functions as a physiological regulator of cargo transport within the axon and our results imply direct effects on the ability of R5L-Tau to modulate motor protein motility. Static Tau complexes are known to modulate the motility of kinesin-1 (KIF5C), acting as “roadblocks” on the microtubule (4, 6, 33, 34, 43, 63, 64). Due to fewer static Tau patches and a reduction in occupancy, we hypothesize R5L-Tau is less inhibitory of kinesin-1 motility compared to WT-Tau. In contrast, diffusive Tau affects the motility of kinesin-3 (KIF1A) (65). Therefore, we would expect R5L-Tau is more inhibitory to KIF1A motility compared to WT-Tau. However, it is also possible that increased flexibility of the projection domain could allow R5L-Tau to directly interact with motor proteins, which has been shown for other MAPs (66, 67). Altogether, the effect of R5L-Tau on cargo transport is an intriguing area of future investigation.

This work has additional implications on the ability of R5L-Tau to regulate microtubules. Previous work has shown, compared to WT-Tau, R5L-Tau reduces total microtubule assembly measured by absorbance at 350 nm (39, 40). The reduction in absorbance could be attributed to a loss of individual microtubule assembly or bundling. The projection domain has a known role in regulating the spacing of microtubule bundles (68), and thus the R5L mutation could reduce Tau induced microtubule bundling due to disruption of the R5 salt bridge with an adjacent microtubule. It is also possible that the R5L mutation reduces the ability of Tau to stabilize individual microtubules due to disruption of patch formation, another interesting avenue of future study.

Finally, our data has implications for pathological Tau. It is proposed that the disease state in many Tauopathies is initiated by promoting Tau aggregation and/or reducing Tau affinity for the microtubule. Aggregation of hyperphosphorylated Tau is a hallmark of Tauopathies, such as PSP, and increased flexibility of the projection domain could alter the phosphorylation state or aggregation propensity of R5L-Tau. There are a number of putative phosphorylation sites in the N-terminal region around the R5 residue (69). However, NMR experiments show no global conformational change in R5L-Tau, and it is therefore unlikely that overall phosphorylation patterns are affected by the mutation. Furthermore, studies of the aggregation propensity of R5L-Tau show no large effect of the mutation (39, 70, 71). In this work, we also show that the R5L mutation, unlike other disease-associated mutations, does not reduce Tau affinity for the microtubule, but alters microtubule bound behavior and disrupts Tau patch formation. Overall, this work challenges the current paradigm of Tauopathies and provides new insight into the role of the N-terminal projection domain in modulating normal and pathological states of Tau on the microtubule surface.

## Materials and Methods

### Purification of Proteins

Tau constructs were purified, characterized, and fluorescently labeled as described (33). Enrichment of ^15^N Tau was performed as described (25). Tubulin was purified from bovine brain (72) and polymerized as described (33). See SI Text for details.

### Microtubule Pelleting Assay

Microtubule pelleting assays were developed based on Charafeddine *et al*., (2019) (73). Briefly, polymerized microtubules and Alexa 488 labeled Tau constructs were incubated for 20 min at 25 °C in pelleting buffer (BRB80 1mM Dithiothreiol (DTT), 10 μg/ml Bovine Serum Albumin (BSA) pH 6.9) with the addition of 10 μM paclitaxel (Sigma Aldrich, St. Louis, MO) for Taxol-microtubules in Bovine Serum Albumin (BSA) coated tubes. The reactions were centrifuged in a TLA-100 rotor in an Optima TLX Ultracentrifuge (Beckman, Pasadena, CA). The supernatant was removed, and the pellet was resuspended in ice cold BRB80 with 10 mg/ml BSA. Fluorescence was measured using a fluorometer (Photon Technology International, Birmingham, NJ) with an excitation of 493 nm and emission of 517 nm. See SI Text for details.

### TIRF Microscopy

Total Internal Reflection Fluorescent (TIRF) microscopy experiments were carried out at room temperature using an inverted Eclipse Ti-E microscope (Nikon, Melville, NY) with 100x Apo TIRF objective lens (1.49 N.A.) and dual iXon Ultra Electron Multiplying CCD cameras, running NIS Elements version 4.51.0. Tau binding assays were completed as previously described with Alexa 488 labeled microtubules and Alexa 647 labeled Tau constructs (43). TIRF dynamics assays using low concentrations of Tau were done as described (43) using Alexa-488 microtubules and Alexa 647 labeled Tau. Three color spiking experiments examining dynamics at high concentrations of Tau were performed using Alexa 405 microtubules, high concentrations of Alexa 488 Tau spiked with 300 pM Alexa 647 Tau. See SI Text for details.

### Circular Dichroism

Circular dichroism spectra of 10 μM WT-Tau and R5L-Tau were acquired on a Chirascan (Applied Photophysics, UK) spectrometer. See SI Text for details.

### NMR Spectroscopy

NMR experiments were performed on a Bruker 700 MHz spectrometer equipped with a cryogenic probe. 2D ^1^H-^15^N HSQC experiments were recorded at 5 °C on 10 μM ^15^N-labelled Tau in BRB80 buffer and 10% D_2_O, in the absence or presence of 20 μM Taxol-microtubules. For the sequence-specific resonance assignment of the peptide (residues 1-20 of Tau), natural abundance 2D ^1^H-^15^N HSQC, 2D ^1^H-^1^H TOCSY and 2D ^1^H-^1^H NOESY were recorded at 5 °C on 2 mM samples in BRB80 buffer and 10% D_2_O. STD experiments were recorded using the Bruker pulse sequence *stdiffgp19.2* with 5 μM Taxol-microtubules and peptides concentrations from 50 μM to 2 mM. See SI Text for details.

## Acknowledgments

We thank David Warshaw and Guy Kennedy for training and support of the TIRF microscope at the University of Vermont. We give special thanks to Vermont Livestock Slaughter & Processing (Ferrisburgh, VT) for supporting our work. This work was supported by National Institute of General Medical Sciences/National Institutes of Health funding to C.L.B. and A.G.H. (GM132646). M.Z. was supported by the Deutsche Forschungsgemeinschaft (DFG, German Research Foundation: SFB 860, TP B2), and the European Research Council (ERC) under the EU Horizon 2020 Research and Innovation Programme (Grant Agreement No. 787679)

## References

1. D. Panda, J. C. Samuel, M. Massie, S. C. Feinstein, L. Wilson, Differential regulation of microtubule dynamics by three- and four-repeat tau: implications for the onset of neurodegenerative disease. Proc Natl Acad Sci U S A 100, 9548–9553 (2003).

2. D. B. Murphy, K. A. Johnson, G. G. Borisy, Role of tubulin-associated proteins in microtubule nucleation and elongation. J Mol Biol 117, 33–52 (1977).

3. G. J. Hoeprich, A. R. Thompson, D. P. McVicker, W. O. Hancock, C. L. Berger, Kinesin’s neck-linker determines its ability to navigate obstacles on the microtubule surface. Biophys J 106, 1691–1700 (2014).

4. D. P. McVicker, L. R. Chrin, C. L. Berger, The nucleotide-binding state of microtubules modulates kinesin processivity and the ability of Tau to inhibit kinesin-mediated transport. J Biol Chem 286, 42873–42880 (2011).

5. M. Vershinin, B. C. Carter, D. S. Razafsky, S. J. King, S. P. Gross, Multiple-motor based transport and its regulation by Tau. Proc Natl Acad Sci U S A 104, 87–92 (2007).

6. R. Dixit, J. L. Ross, Y. E. Goldman, E. L. Holzbaur, Differential regulation of dynein and kinesin motor proteins by tau. Science 319, 1086–1089 (2008).

7. N. M. Kanaan et al., Phosphorylation in the amino terminus of tau prevents inhibition of anterograde axonal transport. Neurobiol Aging 33, 826 e815–830 (2012).

8. N. M. Kanaan et al., Pathogenic forms of tau inhibit kinesin-dependent axonal transport through a mechanism involving activation of axonal phosphotransferases. J Neurosci 31, 9858–9868 (2011).

9. I. Grundke-Iqbal et al., Microtubule-associated protein tau. A component of Alzheimer paired helical filaments. J Biol Chem 261, 6084–6089 (1986).

10. B. Borroni et al., Tau forms in CSF as a reliable biomarker for progressive supranuclear palsy. Neurology 71, 1796–1803 (2008).

11. B. Borroni et al., Tau haplotype influences cerebral perfusion pattern in frontotemporal lobar degeneration and related disorders. Acta Neurol Scand 117, 359–366 (2008).

12. M. Goedert, C. M. Wischik, R. A. Crowther, J. E. Walker, A. Klug, Cloning and sequencing of the cDNA encoding a core protein of the paired helical filament of Alzheimer disease: identification as the microtubule-associated protein tau. Proc Natl Acad Sci U S A 85, 4051–4055 (1988).

13. C. M. Wischik et al., Isolation of a fragment of tau derived from the core of the paired helical filament of Alzheimer disease. Proc Natl Acad Sci U S A 85, 4506–4510 (1988).

14. G. Lindwall, R. D. Cole, Phosphorylation affects the ability of tau protein to promote microtubule assembly. J Biol Chem 259, 5301–5305 (1984).

15. E. M. Mandelkow et al., Tau domains, phosphorylation, and interactions with microtubules. Neurobiol Aging 16, 355–362; discussion 362-353 (1995).

16. B. Zhang et al., Microtubule-binding drugs offset tau sequestration by stabilizing microtubules and reversing fast axonal transport deficits in a tauopathy model. Proc Natl Acad Sci U S A 102, 227–231 (2005).

17. M. Goedert, Tau gene mutations and their effects. Mov Disord 20 Suppl 12, S45–52 (2005).

18. K. H. Strang, T. E. Golde, B. I. Giasson, MAPT mutations, tauopathy, and mechanisms of neurodegeneration. Lab Invest 99, 912–928 (2019).

19. M. D. Weingarten, A. H. Lockwood, S. Y. Hwo, M. W. Kirschner, A protein factor essential for microtubule assembly. Proc Natl Acad Sci U S A 72, 1858–1862 (1975).

20. J. Avila, J. J. Lucas, M. Perez, F. Hernandez, Role of tau protein in both physiological and pathological conditions. Physiol Rev 84, 361–384 (2004).

21. B. L. Goode, M. Chau, P. E. Denis, S. C. Feinstein, Structural and functional differences between 3-repeat and 4-repeat tau isoforms. Implications for normal tau function and the onset of neurodegenetative disease. J Biol Chem 275, 38182–38189 (2000).

22. M. Goedert, M. G. Spillantini, R. Jakes, D. Rutherford, R. A. Crowther, Multiple isoforms of human microtubule-associated protein tau: sequences and localization in neurofibrillary tangles of Alzheimer’s disease. Neuron 3, 519–526 (1989).

23. H. Kadavath et al., Folding of the Tau Protein on Microtubules. Angew Chem Int Ed Engl 54, 10347–10351 (2015).

24. M. D. Mukrasch et al., The “jaws” of the tau-microtubule interaction. J Biol Chem 282, 12230–12239 (2007).

25. H. Kadavath et al., Tau stabilizes microtubules by binding at the interface between tubulin heterodimers. Proc Natl Acad Sci U S A 112, 7501–7506 (2015).

26. B. L. Goode et al., Functional interactions between the proline-rich and repeat regions of tau enhance microtubule binding and assembly. Mol Biol Cell 8, 353–365 (1997).

27. N. Gustke, B. Trinczek, J. Biernat, E. M. Mandelkow, E. Mandelkow, Domains of tau protein and interactions with microtubules. Biochemistry 33, 9511–9522 (1994).

28. B. Trinczek, J. Biernat, K. Baumann, E. M. Mandelkow, E. Mandelkow, Domains of tau protein, differential phosphorylation, and dynamic instability of microtubules. Mol Biol Cell 6, 1887–1902 (1995).

29. E. H. Kellogg et al., Near-atomic model of microtubule-tau interactions. Science 360, 1242–1246 (2018).

30. K. M. McKibben, E. Rhoades, Independent tubulin binding and polymerization by the proline-rich region of Tau is regulated by Tau’s N-terminal domain. J Biol Chem 294, 19381–19394 (2019).

31. A. M. Melo et al., A functional role for intrinsic disorder in the tau-tubulin complex. Proc Natl Acad Sci U S A 113, 14336–14341 (2016).

32. M. H. Hinrichs et al., Tau protein diffuses along the microtubule lattice. J Biol Chem 287, 38559–38568 (2012).

33. D. P. McVicker, G. J. Hoeprich, A. R. Thompson, C. L. Berger, Tau interconverts between diffusive and stable populations on the microtubule surface in an isoform and lattice specific manner. Cytoskeleton (Hoboken) 71, 184–194 (2014).

34. V. Siahaan et al., Kinetically distinct phases of tau on microtubules regulate kinesin motors and severing enzymes. Nat Cell Biol 21, 1086–1092 (2019).

35. R. Tan et al., Microtubules gate tau condensation to spatially regulate microtubule functions. Nat Cell Biol 21, 1078–1085 (2019).

36. M. T. Gyparaki et al., Tau forms oligomeric complexes on microtubules that are distinct from tau aggregates. Proc Natl Acad Sci U S A 118 (2021).

37. A. Deshpande, K. M. Win, J. Busciglio, Tau isoform expression and regulation in human cortical neurons. FASEB J 22, 2357–2367 (2008).

38. P. Poorkaj et al., An R5L tau mutation in a subject with a progressive supranuclear palsy phenotype. Ann Neurol 52, 511–516 (2002).

39. Y. Mutreja, B. Combs, T. C. Gamblin, FTDP-17 Mutations Alter the Aggregation and Microtubule Stabilization Propensity of Tau in an Isoform-Specific Fashion. Biochemistry 58, 742–754 (2019).

40. P. Poorkaj et al., Tau is a candidate gene for chromosome 17 frontotemporal dementia. Ann Neurol 43, 815–825 (1998).

41. B. L. Goode, S. C. Feinstein, Identification of a novel microtubule binding and assembly domain in the developmentally regulated inter-repeat region of tau. J Cell Biol 124, 769–782 (1994).

42. K. A. Butner, M. W. Kirschner, Tau protein binds to microtubules through a flexible array of distributed weak sites. J Cell Biol 115, 717–730 (1991).

43. J. L. Stern, D. V. Lessard, G. J. Hoeprich, G. A. Morfini, C. L. Berger, Phosphoregulation of Tau modulates inhibition of kinesin-1 motility. Mol Biol Cell 28, 1079–1087 (2017).

44. A. R. Duan et al., Interactions between Tau and Different Conformations of Tubulin: Implications for Tau Function and Mechanism. J Mol Biol 429, 1424–1438 (2017).

45. B. T. Castle, K. M. McKibben, E. Rhoades, D. J. Odde, Tau Avoids the GTP Cap at Growing Microtubule Plus-Ends. iScience 23, 101782 (2020).

46. S. M. Kelly, T. J. Jess, N. C. Price, How to study proteins by circular dichroism. Biochim Biophys Acta 1751, 119–139 (2005).

47. S. Kosol, S. Contreras-Martos, C. Cedeno, P. Tompa, Structural characterization of intrinsically disordered proteins by NMR spectroscopy. Molecules 18, 10802–10828 (2013).

48. M. D. Mukrasch et al., Structural polymorphism of 441-residue tau at single residue resolution. PLoS Biol 7, e34 (2009).

49. M. Mayer, B. Meyer, Characterization of Ligand Binding by Saturation Transfer Difference NMR Spectroscopy. Angew Chem Int Ed Engl 38, 1784–1788 (1999).

50. S. Walpole, S. Monaco, R. Nepravishta, J. Angulo, STD NMR as a Technique for Ligand Screening and Structural Studies. Methods Enzymol 615, 423–451 (2019).

51. H. Kadavath et al., The Binding Mode of a Tau Peptide with Tubulin. Angew Chem Int Ed Engl 57, 3246–3250 (2018).

52. S. Barghorn et al., Structure, microtubule interactions, and paired helical filament aggregation by tau mutants of frontotemporal dementias. Biochemistry 39, 11714–11721 (2000).

53. I. D’Souza et al., Missense and silent tau gene mutations cause frontotemporal dementia with parkinsonism-chromosome 17 type, by affecting multiple alternative RNA splicing regulatory elements. Proc Natl Acad Sci U S A 96, 5598–5603 (1999).

54. M. Hong et al., Mutation-specific functional impairments in distinct tau isoforms of hereditary FTDP-17. Science 282, 1914–1917 (1998).

55. M. Hasegawa, M. J. Smith, M. Goedert, Tau proteins with FTDP-17 mutations have a reduced ability to promote microtubule assembly. FEBS Lett 437, 207–210 (1998).

56. P. Rizzu et al., Mutation-dependent aggregation of tau protein and its selective depletion from the soluble fraction in brain of P301L FTDP-17 patients. Hum Mol Genet 9, 3075–3082 (2000).

57. S. Jeganathan, M. von Bergen, H. Brutlach, H. J. Steinhoff, E. Mandelkow, Global hairpin folding of tau in solution. Biochemistry 45, 2283–2293 (2006).

58. M. Schwalbe et al., Predictive atomic resolution descriptions of intrinsically disordered hTau40 and alpha-synuclein in solution from NMR and small angle scattering. Structure 22, 238–249 (2014).

59. M. Schwalbe et al., Structural Impact of Tau Phosphorylation at Threonine 231. Structure 23, 1448–1458 (2015).

60. H. E. Feinstein et al., Oligomerization of the microtubule-associated protein tau is mediated by its N-terminal sequences: implications for normal and pathological tau action. J Neurochem 137, 939–954 (2016).

61. K. J. Rosenberg, J. L. Ross, H. E. Feinstein, S. C. Feinstein, J. Israelachvili, Complementary dimerization of microtubule-associated tau protein: Implications for microtubule bundling and tau-mediated pathogenesis. Proc Natl Acad Sci U S A 105, 7445–7450 (2008).

62. N. Hirokawa, Y. Shiomura, S. Okabe, Tau proteins: the molecular structure and mode of binding on microtubules. J Cell Biol 107, 1449–1459 (1988).

63. L. Balabanian, C. L. Berger, A. G. Hendricks, Acetylated Microtubules Are Preferentially Bundled Leading to Enhanced Kinesin-1 Motility. Biophys J 113, 1551–1560 (2017).

64. A. R. Chaudhary, F. Berger, C. L. Berger, A. G. Hendricks, Tau directs intracellular trafficking by regulating the forces exerted by kinesin and dynein teams. Traffic 19, 111–121 (2018).

65. C. L. Berger, D. V. Lessard, The Microtubule Associated Protein Tau Regulates KIF1A Pausing Behavior and Motility. bioRxiv 10.1101/2021.08.11.455914, 2021.2008.2011.455914 (2021).

66. P. J. Hooikaas et al., MAP7 family proteins regulate kinesin-1 recruitment and activation. J Cell Biol 218, 1298–1318 (2019).

67. B. Y. Monroy et al., Competition between microtubule-associated proteins directs motor transport. Nat Commun 9, 1487 (2018).

68. J. Chen, Y. Kanai, N. J. Cowan, N. Hirokawa, Projection domains of MAP2 and tau determine spacings between microtubules in dendrites and axons. Nature 360, 674–677 (1992).

69. G. Simic et al., Tau Protein Hyperphosphorylation and Aggregation in Alzheimer’s Disease and Other Tauopathies, and Possible Neuroprotective Strategies. Biomolecules 6, 6 (2016).

70. T. C. Gamblin et al., In vitro polymerization of tau protein monitored by laser light scattering: method and application to the study of FTDP-17 mutants. Biochemistry 39, 6136–6144 (2000).

71. E. Chang, S. Kim, H. Yin, H. N. Nagaraja, J. Kuret, Pathogenic missense MAPT mutations differentially modulate tau aggregation propensity at nucleation and extension steps. J Neurochem 107, 1113–1123 (2008).

72. M. Castoldi, A. V. Popov, Purification of brain tubulin through two cycles of polymerization-depolymerization in a high-molarity buffer. Protein Expr Purif 32, 83–88 (2003).

73. R. A. Charafeddine et al., Tau repeat regions contain conserved histidine residues that modulate microtubule-binding in response to changes in pH. J Biol Chem 294, 8779–8790 (2019).

