## Supplemental Methods and Data for "The Pathogenic R5L Mutation Disrupts Formation of Tau Complexes on the Microtubule by Altering Local N-Terminal Structure"

**This PDF file includes:**

- Supplementary text
- Figures S1 to S10
- Legends for Movies S1 to S6
- SI References

**Other supplementary materials for this manuscript include the following:**

- Movies S1 to S6

**Supplementary Information Text**

### Methods

**Tau expression, purification and labeling.** Enrichment of isotopic N15 Tau was performed as described [1]. All Tau constructs were purified and characterized as described previously [2]. Tau constructs were labeled using maleimide chemistry, conjugating Alexa 647 C5 or Alexa 488 C5 Maleimide (Invitrogen Molecular Probes, Carlsbad, CA) to the single cysteine residue. Conjugation of Alexa-488 was done as described previously [3]. For Alexa 647 labeling, Tau protein was incubated in the presence of tenfold molar excess dithiothreitol (DTT) (Sigma-Aldrich. St. Louis, MO) and 4 M Guanidine Hydrogen Chloride (GndHCl) for 10 minutes at 37 °C. Excess DTT was removed through 7MWCO Zeba spin desalting column (Pierce, Rockford, IL). Samples were then incubated with a fivefold molar excess of Alexa 647 C5 Maleimide and incubated at room temperature for 24 hours. Excess free dye was removed by dialysis against BRB80 and confirmed by SDS-PAGE. Dye concentrations were measured using a 640 Spectrophotometer (Beckman, Pasadena, CA) with an extinction coefficient of 270,000 $M^{-1}$ $cm^{-1}$ at 651 nm or 73,000 $M^{-1}$ $cm^{-1}$ at 493 nm for Alexa 647 and Alexa 488, respectively. Protein concentrations were determined through a Modified Lowry Assay (Thermofisher Scientific, Waltham, MA). Labeling efficiencies were determined by taking the ratio of dye concentration to Tau concentration. Tau labeling efficiencies were further assessed using mass spectrometry.

**Quantification of Tau labeling by liquid chromatography-mass spectrometry (LCMS).** In preparation for LCMS, fluorescently labeled Tau was digested to peptides. Aliquots of Tau were solubilized by adding 75 µL 0.1% Rapigest SF surfactant (Waters, Milford, MA) and heating at 50 °C for one hour. Solubilized proteins were then reduced by adding 5 µL of 0.1 M DTT and heating (10 min, 100 °C). Proteins were alkylated by addition of 10.4 µL of 100 mM iodoacetamide (Acros Organics, Fair Lawn, NJ) in 50 mM ammonium bicarbonate and incubating in the dark (30 min, 22 °C). Proteins were cleaved into tryptic peptides by addition of 25 µL of 0.2 µg/µL trypsin (Promega, Madison, WI) in 50 mM ammonium bicarbonate and incubation (18 h, 37 °C). Digested peptide samples were dried by centrifugal evaporation and reconstituted in 100 µL of 7% formic acid in 50 mM ammonium bicarbonate and incubated (1 h, 37 °C) to inactivate trypsin and degrade the Rapigest. Samples were once again evaporated and reconstituted in 100 µL of 0.1 trifluoroacetic acid (TFA) for 1 hour at 37 °C, to ensure cleavage of the Rapigest. After a final evaporation, samples were reconstituted in 100 µL TFA and subjected to 5 minutes of centrifugation at 18,800 RCF (Thermo, Sorvall Legend Micro 21R). A 90-µL aliquot was removed from the supernatant for analysis by LCMS.

Tryptic Tau peptides were separated via high-pressure liquid chromatography and analyzed by mass spectrometry (MS) as previously described [4]. Briefly, a 20-µL aliquot of each sample was injected onto a Waters XSelect HSS T3 column using a Dionex UltiMate 3000 LC system. The effluent was directly infused into a Q Exactive Hybrid Quadrupole-Orbitrap mass spectrometer (Thermo) through an electrospray ionization source. Data were collected in data-dependent MS/MS mode, with the five most abundant ions selected for fragmentation.

To quantify fluorescent labeling, the Thermo .raw files were run through Thermo Proteome Discoverer 2.2 (PD) to identify LCMS peaks and quantify peak areas as previously described [4]. LCMS peak areas for the +4 charge variant of the carbamidomethylated peptide CGSLGNIHHKPGGGQVEVK were extracted manually using the Thermo Xcalibur Qual Browser. This peptide contained the cysteine residue which was labeled with the fluorophore. The degree of labeling was quantified using a mass-balance approach from the loss of the carbamidomethylated peptide, due to fluorescent labeling, as previously described for the quantification of phosphorylation [5, 6]. The five best-ionizing peptides identified and quantified by PD were used as internal reference peptides for the mass-balance calculations.

**Tubulin Purification and Microtubule Polymerization.** Tubulin was purified from bovine brain obtained from Vermont Livestock Slaughter & Processing (Ferrisburgh, VT) as described previously using High Molarity PIPES buffer (1 M PIPES, 20 mM EGTA, 10 mM $MgCl_2$, pH 6.9) [7].

For three color experiments, tubulin was labeled with Alexa 405 carboxylic acid, succinimidyl ester (Invitrogen Molecular Probes, Carlsbad, CA) using methods developed by Hyman et al [8]. Purified tubulin was incubated with Polymerization buffer (BRB80 with 1 M $MgCl_2$, 0.1 M GTP (Sigma-Aldrich) and 50% (v/v) glycerol (Fisher Scientific, Waltham, MA)) for 30 minutes at 37 °C. Polymerized microtubules were pelleted at in high pH cushion buffer (0.1 M NaHEPES pH 8.6 (Sigma-Aldrich) at 60,000 rpm at 37 °C for 45 minutes using an Optima TLX Ultracentrifuge (Beckman, Pasadena, CA). The supernatant was aspirated and washed with warm labeling buffer (0.1 M NaHEPES 1 mM $MgCl_2$, 1 mM EGTA 40% (v/v) glycerol pH 8.6). The pellet was resuspended in warm labeling buffer and incubated with 10 fold molar excess of Alexa 405 for 1 hour at 37 °C. Labeling was quenched using an equal volume of quench buffer (2X BRB80, 100 mM L-Glutamic acid potassium salt monohydrate (K-glutamate; Sigma-Aldrich), 40% (v/v) glycerol and incubated for 5 minutes at 37 °C. The quenched mixture was placed on a warm low cushion buffer (60% (v/v) glycerol, 1X BRB80) and centrifuged at 80,000 rpm for 20 minutes at 37 °C. The supernatant was aspirated and the pellet was rinsed with BRB-80. The pellet was resuspended in Depolymerization Buffer (50 mM K-glutamate, 5 mM $MgCl_2$, pH 7.0) for 30 minutes at 4 °C. Depolymerized tubulin was spun at 80,000 rpm for 10 minutes at 4 °C. The supernatant was allowed to polymerize with Polymerization buffer for 30 minutes at 37 °C. The microtubules were layered onto low cushion buffer and pelleted at 80,000 rpm for 20 minutes at 37 °C. The pellet was resuspended in ice cold BRB80 and incubated for 30 minutes at 4 °C. The depolymerized tubulin was then clarified at 80,000 rpm for 20 minutes at 4 °C. Protein and dye concentrations were determined using a 640 Spectrophotometer (Beckman) with an extinction coefficient 115,000 $M^{-1}$ $cm^{-1}$ at 280 nm for tubulin and an extinction coefficient 35,000 $M^{-1}$ $cm^{-1}$ at 401 nm for Alexa 405.

Microtubules were polymerized as described previously [2]. For NMR experiments, Tubulin was purchased from Cytoskeleton Inc. Taxol-microtubules were polymerized through incubation of tubulin in BRB80 buffer (80 mM PIPES, pH 6.8, 1 mM $MgCl_2$, 1 mM EGTA, 1 mM GTP, 1 mM DTT) at 37 °C for 45 min. Subsequently, an equimolar concentration of paclitaxel was added. Unpolymerized tubulin was removed by ultracentrifugation at 40,000 g for 30 min and removal of the supernatant. The microtubule pellet was resuspended in BRB80 buffer for further experiments.

**Microtubule Pelleting Assay.** Microtubule pelleting assays were developed based on Charafeddine *et al.* (2019) [9]. Centrifuge and Eppendorf tubes were coated with 10 mg/ml BSA overnight and dried to reduce non-specific binding of Tau to labware. Unlabeled microtubules (stabilized with paclitaxel (Sigma-Aldrich) or GMPCPP (Jena Biosciences, Jena, Germany) were polymerized as described earlier and diluted to 1 µM and incubated with Alexa 488 Tau constructs (100 nM Tau with Taxol-microtubules and 300 nM Tau for GMPCPP-microtubules) in pelleting buffer (BRB80 1mM DTT, 10, 10 µg/ml BSA pH 6.9) with the addition of 10 µM paclitaxel for Taxol-microtubules for 20 min at 25 °C. The reactions were centrifuged at 50,000 rpm for 20 minutes at 25 °C in a TLA-100 rotor in an Optima TLX Ultracentrifuge (Beckman, Pasadena, CA) for Taxol-microtubules or 65,000 rpm for 20 minutes at 25 °C in an TLA-100 rotor in Ultracentrifuge for GMPCPP-microtubules. The supernatant was removed, and the pellet was resuspended in ice cold BRB80 with 10 mg/ml BSA. Tubes were placed on ice until fluorescence was measured using a fluorometer (Photon Technology International, Birmingham, NJ) with an excitation of 493 nm and emission of 517 nm. Fraction bound was determined. $F_b = 1 – (F_s/F_{tot})$ where $F_s$ is the fluorescence of the supernatant and $F_{tot}$ is the fluorescence of the supernatant in the absence of microtubules.

**TIRF Microscopy.** All microscopy experiments were done using on glass cover slips prepared as described previously [3]. Experiments were performed in TIRF Assay Buffer (TAB) containing (BRB80, 10 mM DTT, and oxygen scavenger system (0.067 mg/ml glucose oxidase, 0.045 mg/ml catalase, and 5.8 mg/ml glucose; Sigma-Aldrich)). Total Internal Reflection Fluorescent (TIRF) microscopy experiments were carried out at room temperature using an inverted Eclipse Ti-E microscope (Nikon, Melville, NY) with 100x Apo TIRF objective lens (1.49 N.A.) and dual iXon Ultra Electron Multiplying CCD cameras, running NIS Elements version 4.51.0.

**TIRF-based Binding Assay.** Tau binding assays were completed as described [3]. Microtubules labeled with Alexa 488 were excited with a 488 laser (20%), passed though 525/50 filter and imaged for 20 frames (10 frames/sec). Tau, labeled with Alexa 647 was excited with a 640 laser (15%), passed through a 655 Long Pass filter and imaged for 20 frames (10 frames/sec).

**TIRF-based Dynamics Assays.** TIRF based two color single molecule dynamics assays were done as described previously [3, 10]. For two color single molecule studies, Tau constructs were flowed into the chamber at 500 pM and incubated for 3 minutes before imaging. Microtubules labeled with Alexa 488 were excited with a 488 laser (20%), passed though 525/50 filter and imaged for 20 frames (10 frames/sec). Tau, labeled with Alexa 647 was excited with a 640 laser (15%), passed through a 655 Long Pass filter and imaged for 300 frames (10 frames/sec).
For three color spiking experiments, after incubation of microtubules, 100 pM 647-Tau was flowed into the chamber to wash away nonadherent microtubules. Subsequently, a higher concentration of Alexa 488 labeled Tau spiked with 300 pM Alexa 647 labeled Tau was flowed into the chamber and incubated for 3 minutes before imaging. Microtubules labeled with Alexa 405 were excited with a 405 laser (10%), passed through a 450/50 filter and imaged for 20 frames (10 frames/sec) Tau, labeled with Alexa 488 and Alexa 647 were imaged simultaneously using the Illumination Sequence Module (NIS-Elements, Nikon) excited with a 488 laser (5%) and 640 laser (10%), passed through a 525/50 filter and 655 Long Pass filter, respectively. Tau was imaged for 600 frames with 488-Tau (2 frames/sec) and the 647-Tau (10 frames/sec).

**Circular Dichroism.** Circular dichroism spectra of WT-Tau and R5L-Tau were acquired on a Chirascan (Applied Photophysics, UK) spectrometer. Samples were diluted in $ddH_2O$ to reach a final protein concentration of 10 µM. Measurements (3 repeats) were performed in a 0.1 cm light path quartz cuvette at 20 °C with a 1.0 nm bandwidth. Baseline correction was performed subtracting the spectrum of the blank acquired with the same parameter settings. Data are expressed in mean residue ellipticity (MRE) [11]. The deconvolution of secondary structure information contained in the spectra was performed using the Dichroweb online calculation tool [12].

**NMR spectroscopy.** NMR experiments were performed on a Bruker 700 MHz spectrometer equipped with a cryogenic probe. 2D $^{1}H$-$^{15}N$ HSQC experiments were recorded at 5 °C on 10 µM $^{15}N$-labelled WT-Tau and R5L-Tau samples in BRB80 buffer and 10% $D_20$, in the absence or presence of 20 µM Taxol-microtubules (1:2 Tau:tubulin heterodimer ratio). $^{1}H$-$^{15}N$ HSQC spectra were acquired with 2048 and 512 points in the direct and indirect dimensions, respectively, and 32 scans. For the sequence-specific resonance assignment of the WT-Tau peptide and the R5L-Tau peptide (residues 1-20 of Tau), natural abundance 2D $^{1}H$-$^{15}N$ HSQC, 2D $^{1}H$-$^{1}H$ TOCSY and 2D $^{1}H$-$^{1}H$ NOESY (mixing times of 100 and 250 ms) were recorded at 5 °C on 2 mM samples in BRB80 buffer and 10% $D_20$. Spectra were processed using Topspin 3.5pl7 (Bruker) and were further analyzed using the software Sparky [13].

STD NMR samples were prepared in BRB80 buffer. The concentration of Taxol-microtubules was kept constant at 5 µM, while the concentrations of the WT-Tau and R5L-Tau peptides varied from 50 µM to 2 mM. STD experiments were recorded using the Bruker pulse sequence *stdiffgp19.2*. Selective irradiation of Taxol-microtubules was achieved using a train of gaussian-shaped pulses set to 25 ms, at a power level of 46 dB. On-resonance and off-resonance frequencies were set to -1.5 ppm and 60 ppm, respectively. For STD build-up curves, $^{1}H$ 1D STD spectra were acquired at 5 °C using irradiation times of 0.5, 1, 1.5, 2, 3, 5 and 7 seconds. $K_D$ values were determined by the acquisition of $^{1}H$ 1D STD spectra at a fixed concentration of Taxol-microtubules (5 µM) and increasing concentrations of WT-Tau or R5L-Tau peptide. In these measurements, an irradiation time of 2 seconds was used.

For STD analysis, sharp and isolated peaks from the amide and methyl region of $^{1}H$ 1D spectra were selected. For any given proton, the STD effect ($\eta_{STD}$) was calculated using the following equation:

$$\eta_{STD} = \frac{I_0 - I_{STD}}{I_0} \qquad (1)$$

STD build up curves were obtained by plotting $\eta_{STD}$ obtained at each saturation time and by fitting them to the following mono-exponential function

$$\eta_{STD(t_{sat})} = \eta_{STD_{max}} \cdot (1 - \exp(-k_{sat} \cdot t_{sat})) \quad (2)$$

For calculation of $K_D$ values, $\eta_{STD}$ values obtained at increasing peptides concentration were multiplied by the excess of ligand ($\varepsilon$) to provide the STD Amplification Factor ($A_{STD}$) [14]. $K_D$ values were then determined by fitting the $A_{STD}$ values to the following equation:

$$A_{STD} = \frac{(\alpha_{STD} \cdot [L])}{(K_D + [L])} \quad (3)$$

### Data Analysis

**TIRF Dynamics Assays.** Individual Tau binding events were tracked frame to frame using the MTrackJ plugin using ImageJ software (v. 2.0.0, National Institute of Health, Bethesda, MD). Only tracks that started and ended during the duration of imaging were considered had to contain a minimum of 5 frames. Tracks were analyzed in a custom Matlab (The MathWorks, Inc., Natick, MA) script to separate different binding states based on cumulative frequency plots of all displacements across all time intervals within a single event. Events were classified based on the percentage of displacements at bin center 0.4, corresponding to displacement of 0.4 µm. If at least 70 percent of all displacements were more than 0.4 µm, the event was classified as diffusive. Therefore, the percent of events in either binding state was dependent on the percentage of displacements, defined as the classification parameter. Once separated, static:diffusive equilibrium and dwell times for each binding state were determined.

Mean square displacement analysis for diffusive bound events was done using a custom Matlab script, where all squared displacements were determined for all time intervals and averaged. In GraphPad Prism version 8.0, (GraphPad Software, La Jolla, CA) the number of data points were plotted against the time interval and fit to an exponential decay curve equation.

$$Y = (Y_0 - Plateau)e^{-Kx} + Plateau \quad (4)$$

where $Y_0$ is the initial Y value, Plateau is Y value at infinity, and K is the rate constant. The half-life was used to determine the number of points used for fitting of the mean square displacement. The appropriate time intervals and square displacements were used for a mean square displacement analysis.

$$< x^2 > = 2Dt \quad (5)$$

where $<x^2>$ is the mean square displacement, 2 is the numerical constant based on one-dimensional diffusion, D is the diffusion coefficient (µm/sec) and t is time (sec).

**TIRF-based Binding Assays.** Binding assay quantification was done as described [3].

**Tau Patch Analysis.** Patch frequency, fluorescent molecules per patch, and Tau distribution along the microtubule was determined by generating kymographs of Tau bound to microtubules. For patch frequency and number of fluorescent molecules per patch, the normalized average intensity along the microtubule was determined using the plot profile tool in ImageJ. Normalized intensity plots were placed into a custom Matlab script to determine patch frequency and number of molecules per patch. For distribution of Tau along the microtubule, the normalized average intensity was plotted as frequency distributions in GraphPad Prism with values above 1% shown.

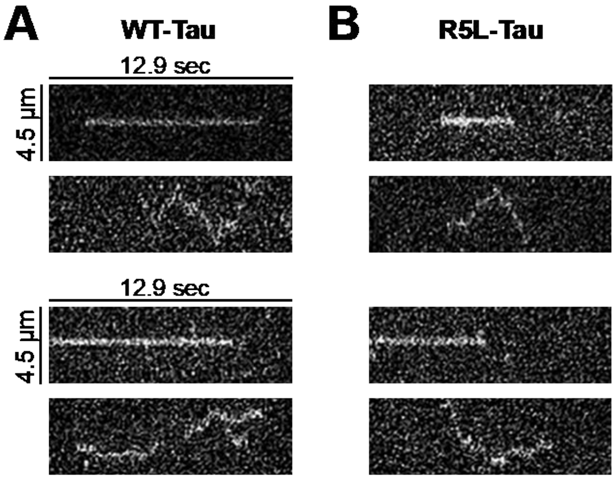


**Fig. S1. Kymographs of WT-Tau and R5L-Tau binding behavior on Taxol-microtubules.**

**A.** Kymographs of WT-Tau (left) and R5L-Tau (right) showing both static (top) and diffusive (bottom) at 500 pM Tau (low concentration). **B.** Kymographs of WT-Tau (left) and R5L-Tau (right) showing both static (top) and diffusive (bottom) at 250 nM Tau (high concentration).

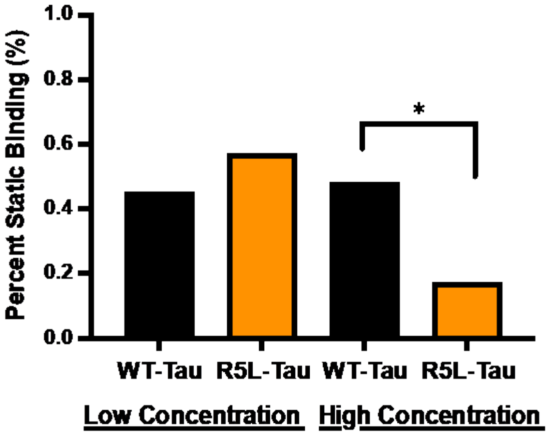


**Fig. S2. N-terminal mutation, R5L, shifts towards diffusive binding at high concentrations on Taxol-microtubules.**

Percent static binding of WT-Tau (black) and R5L-Tau (orange) at either 500 pM Tau (low concentration) or 250 nM Tau (high concentration). For the low concentration, WT-Tau binds statically 45% (N = 439 events) and R5L-Tau 57% (N = 484 events). For the high concentration, WT-Tau binds statically 48% (N = 406 events) while R5L-Tau binding equilibrium shifts towards the diffusive state, binding statically 17% (N = 406 events). Statistical analysis was performed using a Fishers Exact Test (*$p < 0.01$).

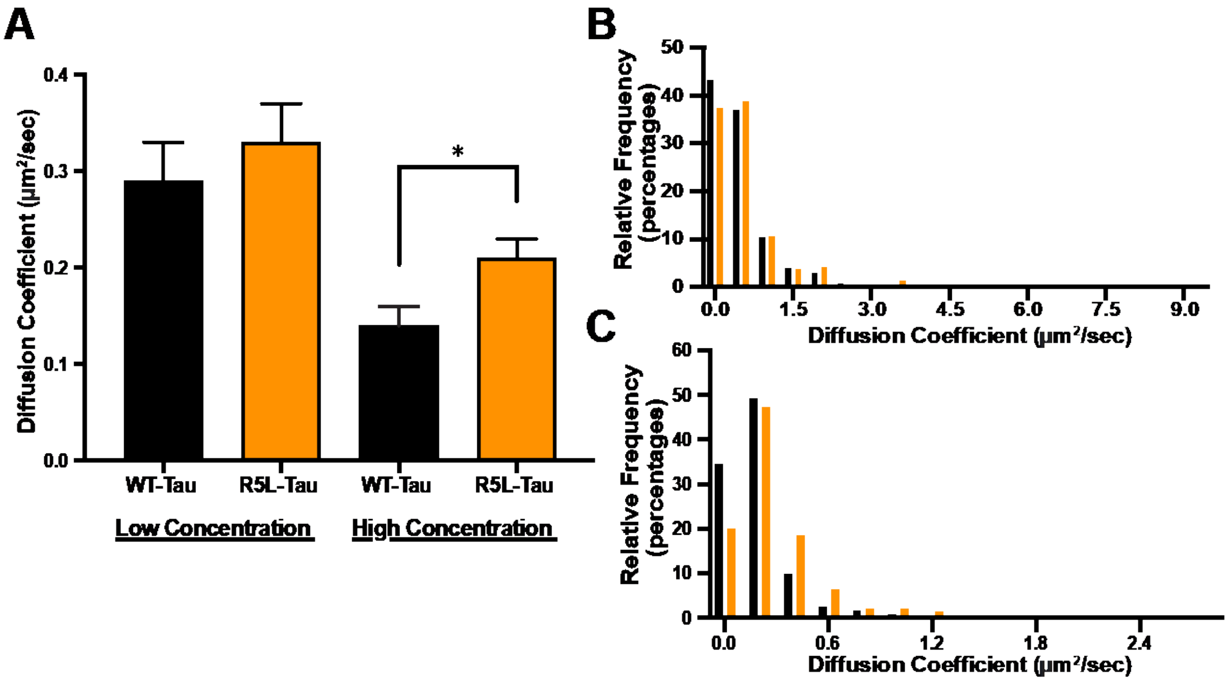


**Fig. S3. WT-Tau diffusion coefficient is reduced at high concentrations on Taxol-microtubules.**

**A.** Diffusion coefficient of WT-Tau (black) and R5L-Tau (orange) at either 500 pM Tau (low concentration) or 250 nM Tau (high concentrations). At the low concentration, WT-Tau had a diffusion coefficient of 0.29 ± 0.04 $\mu m^2$/sec (N= 240 events) and R5L-Tau had a diffusion coefficient of 0.33 ± 0.04 $\mu m^2$/sec (N = 206 events). At the high concentration, WT-Tau had a reduction in diffusion coefficient with a value 0.14 ± 0.2 $\mu m^2$/sec (N = 223 events) compared to R5L-Tau at 0.21 ± 0.2 $\mu m^2$/sec (N = 338 events). Data are median ± 95% CI. Statistical analysis was performed using a Mann-Whitney test. (*$p < 0.01$) **B**. Histograms of diffusion coefficient of WT-Tau (black) and R5L-Tau (orange) at low concentration of Tau. **C.** Histograms of diffusion coefficient of WT-Tau (black) and R5L-Tau (orange) at high concentration of Tau.

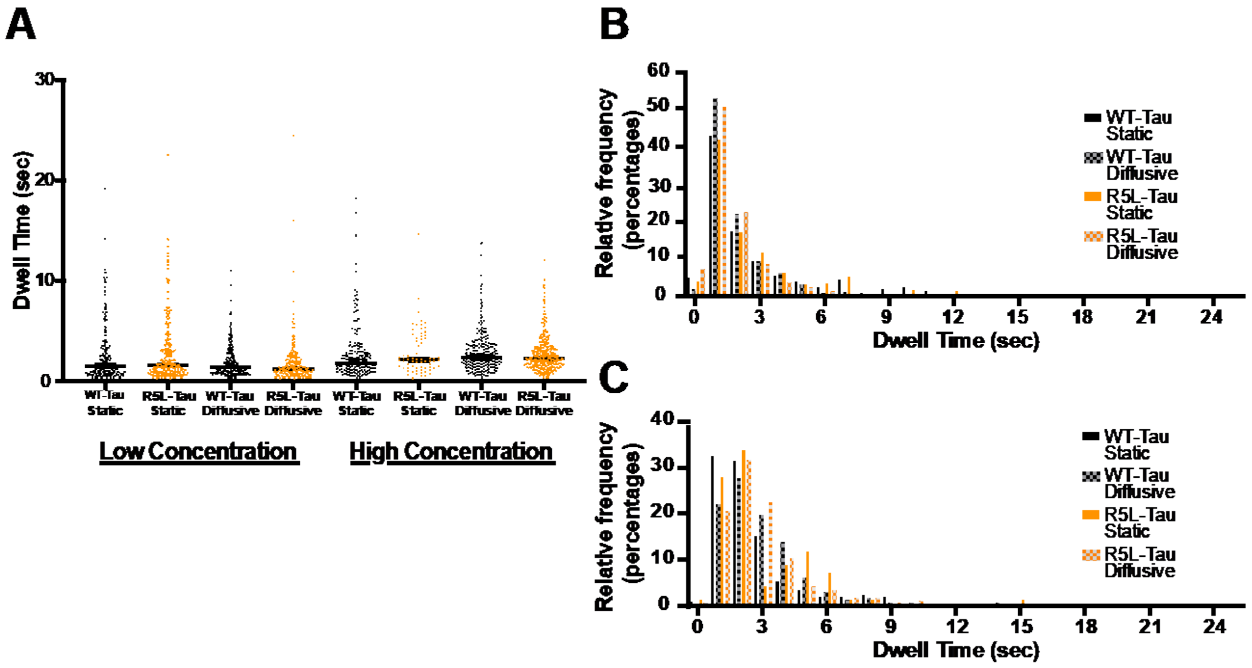


**Fig. S4. Dwell times of WT-Tau and R5L-Tau Taxol-microtubules.**

Dwell times of WT-Tau (black) and R5L-Tau (orange) at either 500 pM Tau (low concentration) or 250 nM Tau (high concentration). **A.** At the low concentration, WT-Tau static binding events had a dwell time of 1.5 ± 0.3 sec (N = 199 events) while diffusive events had a dwell time of 1.4 ± 0.1 sec (N = 240 events). Similarly, R5L-Tau static events had a dwell time of 1.6 ± 0.2 sec (N = 278 events) and diffusive events had a dwell time of 1.2 ± 0.2 sec (N = 206 events). At the high concentration, WT-Tau static events had a dwell time of 1.8 ± 0.4 sec (N = 183 events) and diffusive events bound 2.4 ± 0.3 sec (N = 223 events). R5L-Tau static events bound 2.1 ± 0.3 sec (N = 68 events) and diffusive events 2.3 ± 0.2 sec (N = 338 events). Data are median ± 95% CI. Statistical analysis was performed using a Mann-Whitney test. (*p < 0.01) **B**. Histograms of dwell times of WT-Tau (black) and R5L-Tau (orange) at low concentration of Tau. **C.** Histograms of dwell times of WT-Tau (black) and R5L-Tau (orange) at high concentration of Tau.

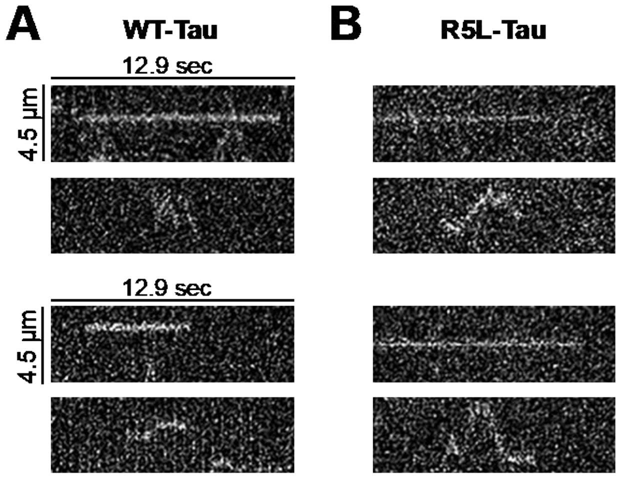


**Fig. S5. Kymographs of WT-Tau and R5L-Tau on GMPCPP-microtubules.**

**A.** Kymographs of WT-Tau (left) and R5L-Tau (right) showing both static (top) and diffusive (bottom) at 500 pM (low concentration). **B.** Kymographs of WT-Tau (left) and R5L-Tau (right) showing both static (top) and diffusive (bottom) at 400 nM (high concentration).

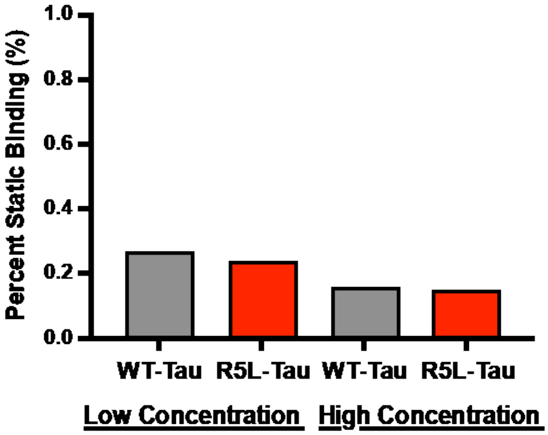


**Fig. S6. WT-Tau and R5L-Tau binding on GMPCPP-microtubules.**

Percent static binding of WT-Tau (grey) and R5L-Tau (red) at either 500 pM Tau (low concentration) or 400 nM Tau (high concentration). At the low concentration, WT-Tau binds statically 26% (N = 424 events) and R5L-Tau 23% (N = 446 events). At the high concentration, WT-Tau binds statically 15% (N = 236 events) and R5L-Tau binds statically 14% (N = 283 events). Statistical analysis was performed using a Fishers Exact Test (*$p < 0.01$).

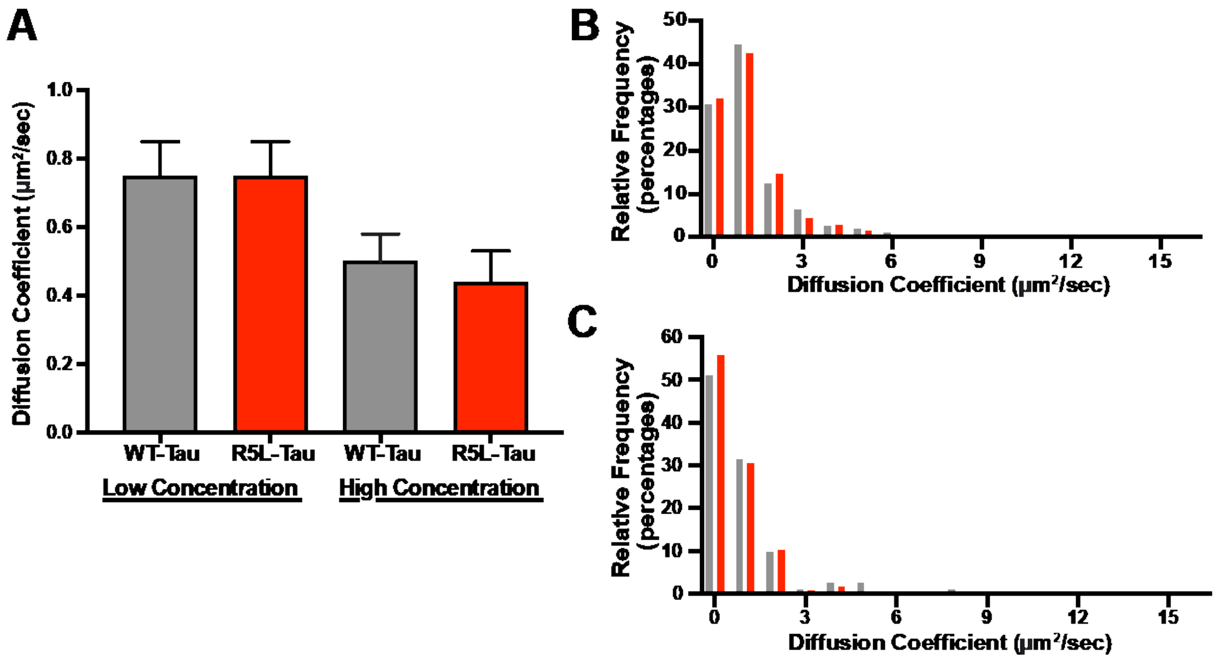


**Fig. S7. Diffusion coefficient of WT-Tau and R5L-Tau on GMPCPP-microtubules.**

**A.** Diffusion coefficient of WT-Tau (grey) and R5L-Tau (red) at either 500 pM Tau (low concentration) or 400nM Tau (high concentration). At the low concentration, WT-Tau had a diffusion coefficient of 0.75 ± 0.10 $\mu m^2$/sec (N = 313 events) and R5L-Tau had a diffusion coefficient of 0.75 ± 0.10 $\mu m^2$/sec (N = 342 events). At high concentrations, WT-Tau had a diffusion coefficient of 0.50 ± 0.08 $\mu m^2$/sec (N= 200 events) and R5L-Tau 0.44 ± 0.09 $\mu m^2$/sec (N = 242 events). Data are median ± 95% CI. Statistical analysis was performed using a Mann-Whitney test. (*$p < 0.01$) B. Histograms of diffusion coefficient of WT-Tau (grey) and R5L-Tau (red) at low concentration of Tau. **C.** Histograms of diffusion coefficient of WT-Tau (grey) and R5L-Tau (red) at high concentration of Tau.

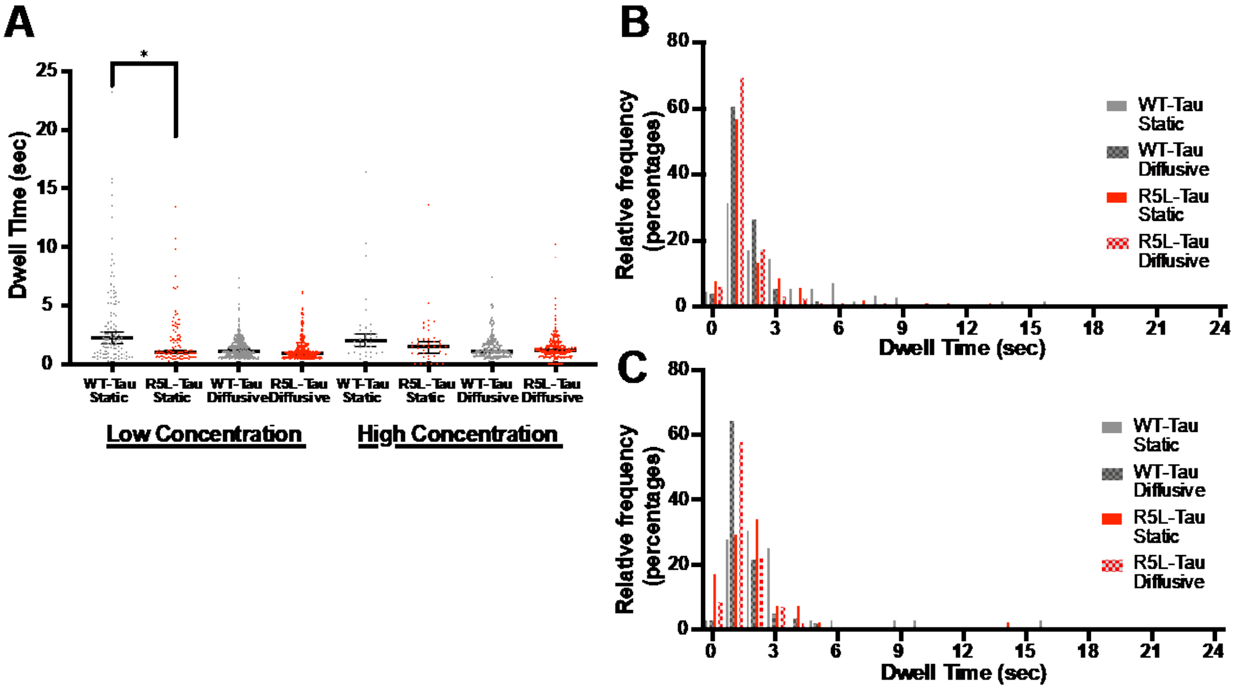


**Fig. S8. Dwell times of WT-Tau and R5L-Tau on GMPCPP-microtubules.**

Dwell times of WT-Tau (grey) and R5L-Tau (red) at either 500 pM Tau (low concentration) or 400 nM Tau (high concentration). **A.** At the low concentration, WT-Tau static events had a dwell time of 2.2 ± 0.5 sec (N = 110 events) and diffusive events 1.1 ± 0.2 sec (N = 313 events). For R5L-Tau, static events had a dwell time of 1.0 ± 0.2 sec (N = 107 events) and diffusive events 0.9 ± 0.1 sec (N = 342 events). At the high concentration, WT-Tau static events had a dwell time of 2.0 ± 0.6 sec (N = 36 events) and diffusive event dwell time of 1.5 ± 0.4 sec (N = 200 events). R5L-Tau static events had a dwell time of 1.1 ± 0.1 sec (N = 40 events) and diffusive events had a dwell time of 1.2 ± 0.1 sec (N = 242 events). Data are median ± 95% CI. Statistical analysis was performed using a Mann-Whitney test. (*$p < 0.01$) **B.** Histograms of dwell times of WT-Tau (grey) and R5L-Tau (red) at low concentration of Tau. **C.** Histograms of dwell times of WT-Tau (grey) and R5L-Tau (red) at high concentration of Tau.

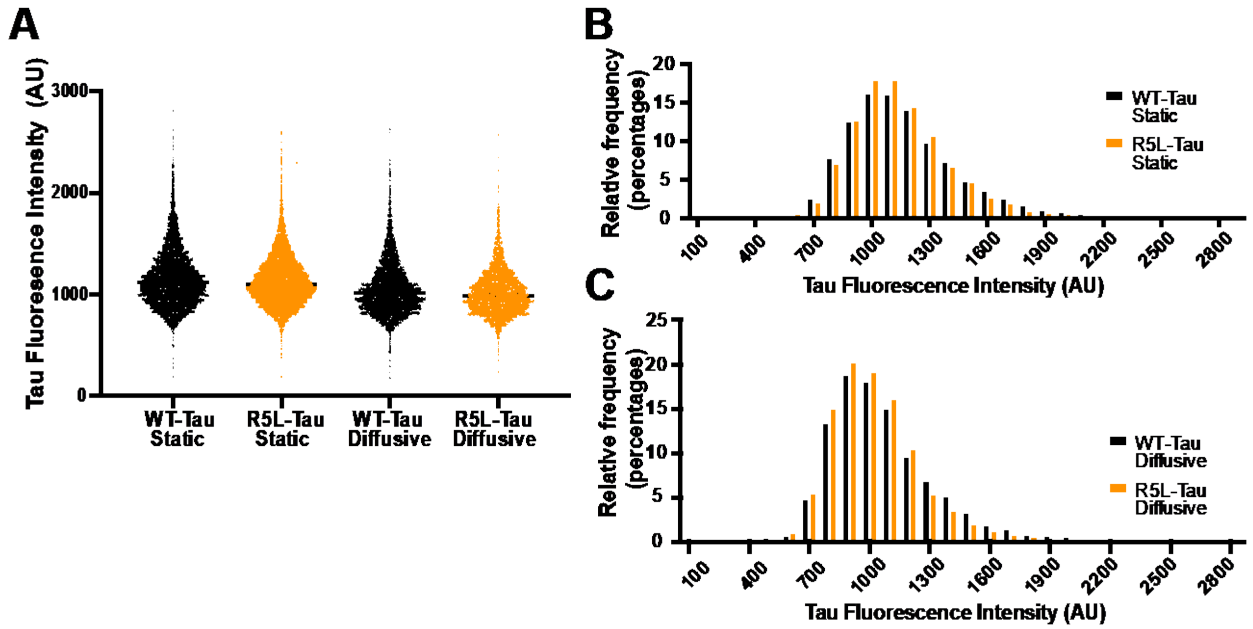


**Fig. S9. Fluorescence Intensity of individual WT-Tau and R5L-Tau on Taxol-microtubules.**

**A.** Fluorescence intensity of individual frames of WT-Tau (black) and R5L-Tau (orange) Alexa 647 labeled Tau bound to Taxol-microtubules showing both static and diffusive events. The fluorescence intensity of static events off WT-Tau 1116 ± 9 (N = 5628 frames) and R5L-Tau 1103 ± 7 (N = 8028 frames). The fluorescence intensity of diffusive events WT-Tau 1017 ± 10 (N = 4736 frames) and R5L-Tau 991 ± 10 (N = 3769 frames) Data are median ± 95% CI **B**. Histograms of fluorescence intensity of static events. **C.** Histograms of fluorescence intensity of diffusive events. Statistical analysis was performed using a Mann-Whitney test (*p < 0.01)

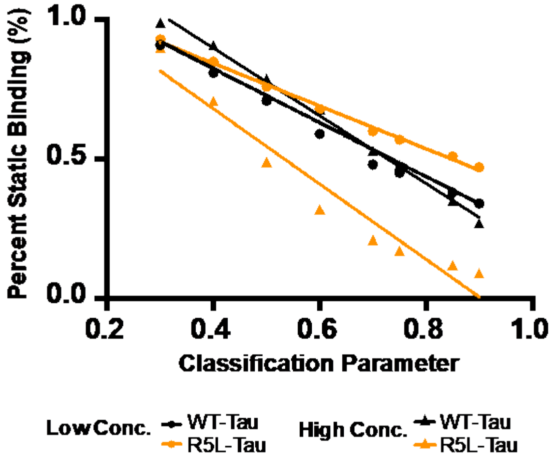


**Fig. S10. Percent static molecules is dependent on threshold for static events.**
The percent of static molecules of WT-Tau (black) or R5L-Tau (orange) at 500 pM Tau (low concentration, circles) or 250 nM Tau (high concentration, triangles) on Taxol-microtubules are plotted against the threshold of static events. At the low concentration, WT-Tau and R5L-Tau have similar slopes and converge on the same point. At the high concentration, WT-Tau has a higher value than R5L-Tau at each data point. Comparing low and high concentrations for either WT-Tau or R5L-Tau, there is a larger slope at the high concentration compared to the low concentration of Tau.

**Movie S1**. WT-Tau binding to Taxol-Microtubules. Representative movie of 250 nM Alexa 647 labeled WT-Tau on Taxol-microtubules imaged at 10 frames/sec. Scale bar = 2 μm.

**Movie S2**. R5L-Tau binding to Taxol-Microtubules. Representative movie of 250 nM Alexa 647 labeled R5L-Tau on Taxol-microtubules imaged at 10 frames/sec. Scale bar = 2 μm.

**Movie S3**. WT-Tau behavioral dynamics at high concentrations on Taxol-microtubules. 250 nM Alexa 488 labeled Tau (green) spiked with 300 pM Alexa 647 labeled Tau (pink) on Alexa 405 labeled Taxol-microtubules (blue). Scale bar = 2 μm. Individual molecules were imaged at 10 frames/sec.

**Movie S4**. R5L-Tau patches behavioral dynamics at high concentrations on Taxol-microtubules. 250 nM Alexa 488 labeled Tau (green) spiked with 300 pM Alexa 647 labeled Tau (pink) on Alexa 405 labeled Taxol-microtubules (blue). Scale bar = 2 μm. Individual molecules were imaged at 10 frames/sec.

**Movie S5**. WT-Tau binding to GMPCPP-microtubules. Representative movie of 450 nM Alexa 647 labeled WT-Tau on GMPCPP-microtubules imaged at 10 frames/sec. Scale bar = 2 μm.

**Movie S6**. WT-Tau binding to GMPCPP-microtubules. Representative movie of 450 nM Alexa 647 labeled R5L-Tau on GMPCPP-microtubules imaged at 10 frames/sec. Scale bar = 2 μm.

**Si References**


1. Kadavath, H., et al., *Tau stabilizes microtubules by binding at the interface between tubulin heterodimers.* Proc Natl Acad Sci U S A, 2015. **112**(24): p. 7501-6.
2. McVicker, D.P., L.R. Chrin, and C.L. Berger, *The nucleotide-binding state of microtubules modulates kinesin processivity and the ability of Tau to inhibit kinesin-mediated transport.* J Biol Chem, 2011. **286**(50): p. 42873-80.
3. Stern, J.L., et al., *Phosphoregulation of Tau modulates inhibition of kinesin-1 motility.* Mol Biol Cell, 2017. **28**(8): p. 1079-1087.
4. O'Leary, T.S., et al., *MYBPC3 truncation mutations enhance actomyosin contractile mechanics in human hypertrophic cardiomyopathy.* J Mol Cell Cardiol, 2019. **127**: p. 165-173.
5. Previs, M.J., et al., *Quantification of protein phosphorylation by liquid chromatography-mass spectrometry.* Anal Chem, 2008. **80**(15): p. 5864-72.
6. Previs, M.J., et al., *Molecular mechanics of cardiac myosin-binding protein C in native thick filaments.* Science, 2012. **337**(6099): p. 1215-8.
7. Castoldi, M. and A.V. Popov, *Purification of brain tubulin through two cycles of polymerization-depolymerization in a high-molarity buffer.* Protein Expr Purif, 2003. **32**(1): p. 83-8.
8. Hyman, A., et al., *Preparation of modified tubulins.* Methods Enzymol, 1991. **196**: p. 478-85.
9. Charafeddine, R.A., et al., *Tau repeat regions contain conserved histidine residues that modulate microtubule-binding in response to changes in pH.* J Biol Chem, 2019. **294**(22): p. 8779-8790.
10. McVicker, D.P., et al., *Tau interconverts between diffusive and stable populations on the microtubule surface in an isoform and lattice specific manner.* Cytoskeleton (Hoboken), 2014. **71**(3): p. 184-94.
11. Kelly, S.M., T.J. Jess, and N.C. Price, *How to study proteins by circular dichroism.* Biochim Biophys Acta, 2005. **1751**(2): p. 119-39.
12. Whitmore, L. and B.A. Wallace, *Protein secondary structure analyses from circular dichroism spectroscopy: methods and reference databases.* Biopolymers, 2008. **89**(5): p. 392-400.
13. Lee, W., M. Tonelli, and J.L. Markley, *NMRFAM-SPARKY: enhanced software for biomolecular NMR spectroscopy.* Bioinformatics, 2015. **31**(8): p. 1325-7.
14. Mayer, M. and B. Meyer, *Group epitope mapping by saturation transfer difference NMR to identify segments of a ligand in direct contact with a protein receptor.* J Am Chem Soc, 2001. **123**(25): p. 6108-17.
